# Oncogenic Wnt/STOP signaling regulates ribosome biogenesis *in vivo*

**DOI:** 10.1101/326819

**Authors:** Babita Madan, Nathan Harmston, Gahyathiri Nallan, Alex Montoya, Peter Faull, Enrico Petretto, David M. Virshup

**Author notes:** BM and NH contributed equally to this work. Lead Contact: David M. Virshup.

## Abstract

Activating mutations in the Wnt pathway drive a variety of cancers, but the specific targets and pathways activated by Wnt ligands are not fully understood. To bridge this knowledge gap, we performed a comprehensive time-course analysis of Wnt-dependent signaling pathways in an orthotopic model of Wnt-addicted pancreatic cancer, using a PORCN inhibitor currently in clinical trials, and validated key results in additional Wnt-addicted models. The analysis of temporal changes following Wnt withdrawal demonstrated direct and indirect regulation of >3,500 Wnt activated genes (23% of the transcriptome). Regulation was both transcriptional via Wnt/β-catenin, and through the modulation of protein abundance of important transcription factors including MYC via Wnt/STOP. Our study identifies a central role of Wnt /β-catenin and Wnt/STOP signaling in controlling ribosomal biogenesis, a key driver of cancer proliferation.

## Introduction

Wnts are a family of 19 secreted proteins that play key roles in cell proliferation, cell-cell communication and differentiation and are essential during embryonic development and in adult tissue homeostasis (Nusse and Clevers, 2017). The binding of Wnts to their receptors and co-receptors results in the regulation of multiple downstream signaling pathways (Acebron and Niehrs, 2016). Our knowledge of the specific events and signals regulated by Wnts derives from a variety of genetic, molecular, and biochemical approaches that has generated a rich map of these downstream pathways (Ramakrishnan and Cadigan, 2017; Yu, 2014).

The Wnt/β-catenin pathway, also known as canonical WNT signaling, has been intensively studied. In the presence of Wnts, β-catenin is stabilized and translocates to the nucleus where it drives expression of target genes in a context specific manner via binding to TCF7L2 and other factors (Ju et al., 2014; Nakamura et al., 2016; Ramakrishnan and Cadigan, 2017). In addition to the Wnt/β-catenin pathway, Wnts regulate signaling through diverse β-catenin independent non-canonical pathways such as the PCP (planar cell polarity) pathway and the Wnt/STOP (Wnt-dependent stabilization of proteins) pathway that are less well characterized (Acebron et al., 2014; Chien et al., 2009; Koch et al., 2015; Taelman et al., 2010; Zhang et al., 2012).

While the downstream mutations that stabilize β-catenin (e.g. in the adenomatous polyposis coli (*APC*) gene) clearly cause human cancers, genetic lesions that cause Wnt over-expression have not been found (Nusse and Varmus, 2012). A subset of mutations that block Wnt receptor internalization and confer dependency on Wnt ligands have been identified in a range of carcinomas. These include loss of function mutations in RNF43, an E3-ligase, and translocations leading to increased R-spondin levels (Jiang et al., 2013; Ong et al., 2012; Seshagiri et al., 2012). RNF43 mutations(Cancer Genome Atlas Research Network, 2017) and translocations involving RSPO2 and RSPO3 are found in 7% of pancreatic adenocarcinoma (PDAC) and 10% of colorectal cancers respectively (Seshagiri et al., 2012). Cancers with RNF43 or RSPO3 mutations have a markedly increased abundance of Frizzled receptors on the cell surface and are uniquely Wnt-addicted (Madan et al., 2016; Madan and Virshup, 2015).

Wnts are palmitoleated by a membrane bound O-acyltransferase, PORCN. This modification is(Rios-Esteves and Resh, 2013) essential for binding to chaperone WLS and Frizzled receptors, and is therefore required for the activity of all Wnts. Pharmacological PORCN inhibitors such as ETC-159 and LGK-974 have progressed to Phase I clinical trials due to their efficacy in preclinical models of RNF43-mutant pancreatic and RSPO3-translocated colorectal cancers (Jiang et al., 2013; Madan et al., 2016; Proffitt et al., 2013). The recent development of these PORCN inhibitors that block all Wnt secretion provides an opportunity to investigate how Wnt regulated genes change over time following withdrawal of signaling (Chen et al., 2009; Coombs et al., 2010; Janda et al., 2012; Liu et al., 2013; Madan et al., 2016; Takada et al., 2006). In order to provide the most relevant data, it is important to use the most predictive pre-clinical models. A large body of literature demonstrates that cancers *in vivo* behave very differently than cancers in tissue culture (Killion et al., 1998). These differences have led to the development of orthotopic xenograft models and patient-derived xenografts that better reflect the behavior of Wnt-addicted cancers in a complex cancer-host environment (Byrne et al., 2017).

Here we investigated the temporal impact of acute withdrawal of Wnt ligands on the perturbed transcriptome of a Wnt-addicted human pancreatic cancer in an orthotopic mouse model. The time-series analysis identified direct and indirect Wnt targets based on their dynamics, distinguishing immediate early, early and late response genes. This analysis led to identification of an important role of the Wnt/STOP pathway in regulating tumor growth. We further disentangled the WNT *vs* MYC dependencies in these cancers. This comprehensive time course analysis in an *in vivo* Wnt-addicted cancer provides both a valuable resource and new insights into the central role of Wnt/β-catenin and Wnt/STOP signaling in regulation of ribosomal biogenesis pathway.

## Results

### Time-dependent global transcriptional changes follow PORCN inhibition in a Wnt-addicted pancreatic cancer model

We aimed to identify the genes and biological processes that are directly or indirectly regulated by Wnt ligands in a Wnt-addicted cancer *in vivo*. To mirror the tumor microenvironment and recapitulate tumor stromal interactions, we established an orthotopic mouse model using a highly WNT-dependent HPAF-II cell line with *RNF43* inactivating mutation (Jiang et al., 2013; Madan et al., 2016). As expected, ETC-159 significantly inhibited the growth of HPAF-II orthotopic xenografts (Fig. 1A), and led to pronounced histomorphological changes (Fig. 1B) (Madan et al., 2016). Tumors from the control group were characterized by the presence of neoplastic cells with poorly defined acini and cell boundaries. Treatment with ETC-159 induced changes in cellular organization and by 7 days the tissue appeared more differentiated with a decreased nuclear cytoplasmic ratio, and diminished anisocytosis and anisokaryosis.

**Figure 1:**
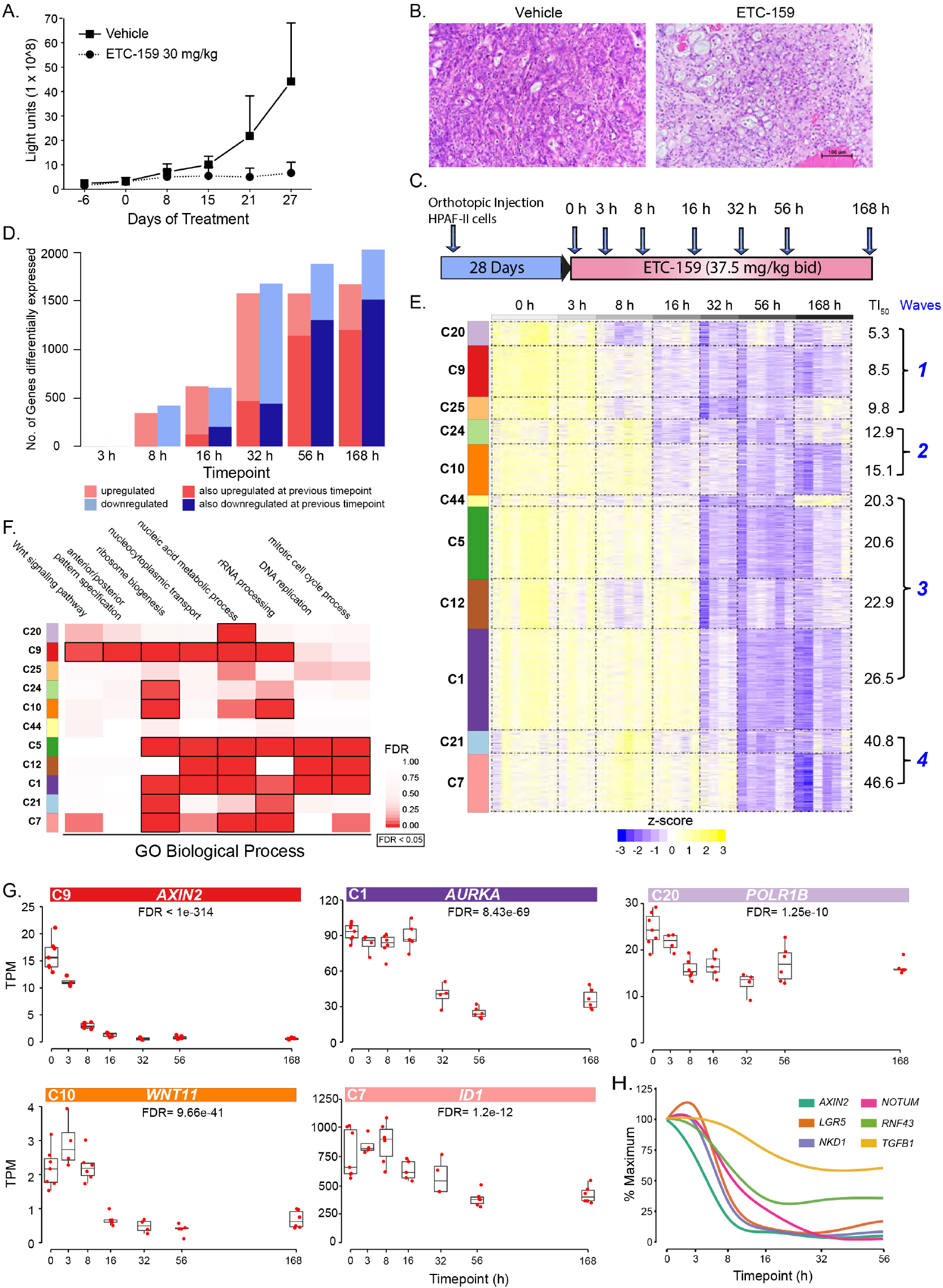
PORCN inhibition remodels the transcriptome of RNF43 mutant pancreatic cancer. A. ETC-159 treatment prevents the growth of orthotopic HPAF-II xenografts. HPAF-II cells (10^6^) were injected into the tail of the mouse pancreas. Following establishment of tumors (~3 weeks), the mice were treated daily with 30 mg/kg ETC-159. Tumorgrowth was monitored by measuring luciferase activity. Data represent mean +/− standard deviation (SD). N = 8/group.
B. Inhibition of Wnt signaling promotes histological changes in HPAF-II xenografts. Hematoxylin & eosin stained images of HPAF-II xenografts treated with ETC-159 for 28 days.
C. Schematic representation of the Experimental plan. HPAF-II cells (10^6^) were injected into the tail of the mouse pancreas. Following establishment of tumors (28 d), the mice were treated twice daily with 37.5 mg/kg/dose ETC-159. Tumors were harvested at the indicated time points.
D. ETC-159 treatment leads to widespread changes in the transcriptome. Total number of genes whose expression changes after PORCN inhibition over time compared to 0 h. Genes whose expression was up or downregulated at the previous time point are also indicated (absolute fold change > 1.5, FDR < 10%).
E. Time series clustering of *Wnt-activated* genes reveals distinct patterns of response to PORCN inhibition. Genes differentially expressed over time (FDR < 10%) in response to PORCN inhibition were clustered by expression pattern using GPClust. The cluster number (C) and TI_50_ (time to a 50% decrease in gene expression) for each cluster is indicated.
F. GO Biological Process enrichments of each cluster of *Wnt-activated* genes. Enrichment of *Wnt-activated* genes highlights processes including Wnt signaling, ribosome biogenesis and the cell cycle.
G. Illustration of the various time courses of gene expression following PORCN inhibition. Representative *Wnt-activated* genes from selected clusters are shown. (TPM, transcripts per million reads)
H. Well-established Wnt/β-catenin target genes change with distinct dynamics following PORCN inhibition.

To identify immediate early, early and late responses to Wnt inhibition, mice with established HPAF-II orthotopic xenografts were treated with ETC-159 and tumors were collected at 3, 8, 16, 32, 56 hours and at 7 days after treatment (Fig. 1C). Comprehensive gene expression analysis was performed using RNA-seq of 4-7 independent tumors at each time point (Fig. S1A). Inhibition of WNT signaling led to a marked change in the transcriptome, with the expression of 11,673 genes (75% of all expressed genes) changing over time (false discovery rate (FDR) < 10%) (Table S1). Expression of 773 genes changed as early as 8 h after the first dose of ETC-159. 1,578 and 1,883 genes were upregulated or downregulated, respectively after 56 h (FDR < 10%, absolute fold-change > 1.5) (Figs 1D, S1B-C). The majority of genes that exhibited significant differences at 56 h were also differentially expressed at 7 days, suggesting that the effect of Wnt inhibition is primarily established within 3 days.

To better understand how the withdrawal of Wnt signaling affected gene expression over time we performed time-series clustering (Hensman et al., 2015) of differentially expressed genes. Genes with significant changes in response to treatment were grouped into 64 clusters, with each cluster consisting of genes exhibiting similar dynamic responses following PORCN inhibition (Fig. S1D, Table S2). Further analysis of these clusters to identify consistent global patterns of transcriptional response identified two major robust patterns (Fig. S1E-F), a supercluster comprising of genes consistently down-regulated (*Wnt-activated genes*) and a supercluster containing genes consistently upregulated following PORCN inhibition (*Wnt-repressed genes*) (Fig. 1E). Here, to demonstrate the usefulness of this data resource, we focus on *Wnt-activated* genes, and we provide the analysis of Wnt-ligand regulated genes in Tables S1-S2.

### Analysis of Wnt-activated genes

The *Wnt-activated* genes supercluster contained 11 clusters with distinct dynamics, consisting of 3,549 genes (23% of the transcriptome). For each of these clusters, we calculated the time to 50% inhibition (TI_50_) based on its mean profile. We operatively classified these clusters into four waves with TI_50_ ranging from 5.3 to 46.6 hours (Fig. 1E) (first wave, 5-3-9.8 h; second, 12.9-15.1; third, 20.3-26.5; fourth, 40.8-46.6). Well-established Wnt target genes had distinct time courses and were present throughout the first three waves with TI_50_ ranging from 5.3 to 26.5 hrs. For example, Cluster 9 (TI_50_ = 8.5 h) included well-known β-catenin targets (e.g. *AXIN2*, *NKD1*, *RNF43*, *BMP4* and *LGR5*) and was significantly enriched for pathways and processes relating to Wnt signaling and development (Fig. 1F). C5 and C12 in the third wave (TI_50_ 20.6 and 22.9) similarly contained known Wnt target genes *e.g. NOTUM* (Giraldez et al., 2002). We speculate that these complex dynamics and broad range of response times of the various β-catenin targets relates to the cell-type specific context of the co-regulatory elements of these genes and the stability of the specific mRNAs.

Interestingly, we observed that the early-changing clusters such as C9 contained genes that are not known to be direct β-catenin target genes (Fig. S2) and thus may be β-catenin independent and rely on mechanisms such as Wnt/STOP. These included well-studied regulators of ribosomal biogenesis (e.g. *NPM1*, *DKC1*, *NOL6*, *RRS1*) and nucleocytoplasmic transport (e.g. *XPO5*, *NUP37*) (van Riggelen et al., 2010). Other smaller rapidly responding clusters in the first and second waves similarly contained genes not known to be β-catenin target genes. These clusters were also enriched for processes associated with ribosome biogenesis (e.g. C20: *POLR1A*, *POLR1B*; C25: *NOP14*, *RRP9*). The slowest responding genes, found in the fourth wave clusters 7 and 21 (TI_50_ 40.8-46.6 h) are likely to be regulated by processes downstream of initial Wnt signaling events and were also enriched for processes relating to ribosomal biogenesis (Fig. 1E & F). In addition to ribosome biogenesis, there was a broad enrichment throughout the clusters for genes involved in nucleic acid metabolism and cell cycle, especially in the third wave, C1, C5 and C12 (Fig. 1F). Selected examples of *Wnt-activated* genes with differences in their pattern of response to Wnt inhibition are depicted in Fig. 1G.

Our dataset provides a comprehensive resource of genes whose expression is highly dependent on Wnt signaling *in vivo* (Table S1). As several of the early changing genes are not known to be direct targets of β-catenin, this analysis identified genes that may depend on additional pathways such as Wnt/STOP. Importantly, the dataset highlights that in addition to its recognized role in cell cycle regulation, an early consequence of blocking Wnt signaling is the down-regulation of genes involved in ribosome biogenesis and its associated processes.

### A common core of Wnt-regulated gene expression changes are more robust in orthotopic xenografts

To test if the gene expression changes seen in the HPAF-II pancreatic cancer were generalizable to other Wnt-addicted cancers, we compared our data to our previously published dataset of a Wnt-addicted RSPO3-translocation patient-derived colorectal cancer xenograft (CRC PDX) (Table S3) (Madan et al., 2016). The strength of the correlation between the expression changes induced by PORCN inhibition at 56 h in the two experimental systems (r^2^ = 0.36, Fig. 2A) indicated that regardless of the upstream mutation and tissue of origin, the downstream effect of Wnt inhibition on tumor gene expression was similar. In keeping with the central role of Wnts in regulating the differentially expressed genes, the majority of *Wnt-activated* genes (62%, FDR < 10%, absolute fold-change > 1.5) were also down regulated at 56 h in the CRC PDX (Fig. 2B). Within C9, 69% of the genes expressed in both systems were down-regulated in CRC PDX, suggesting that these genes may be direct targets of WNT signaling in both of these tumor models. This highlights that the core processes and mechanisms responsible for the *Wnt-activated* genes are shared between the CRC PDX and HPAF-II orthotopic xenografts. Notably, the overlapping set of *Wnt-activated* genes was enriched for genes involved in cell cycle regulation and ribosome biogenesis (Fig. S3A), again suggesting the centrality of these pathways in Wnt-addicted cancers.

**Figure 2:**
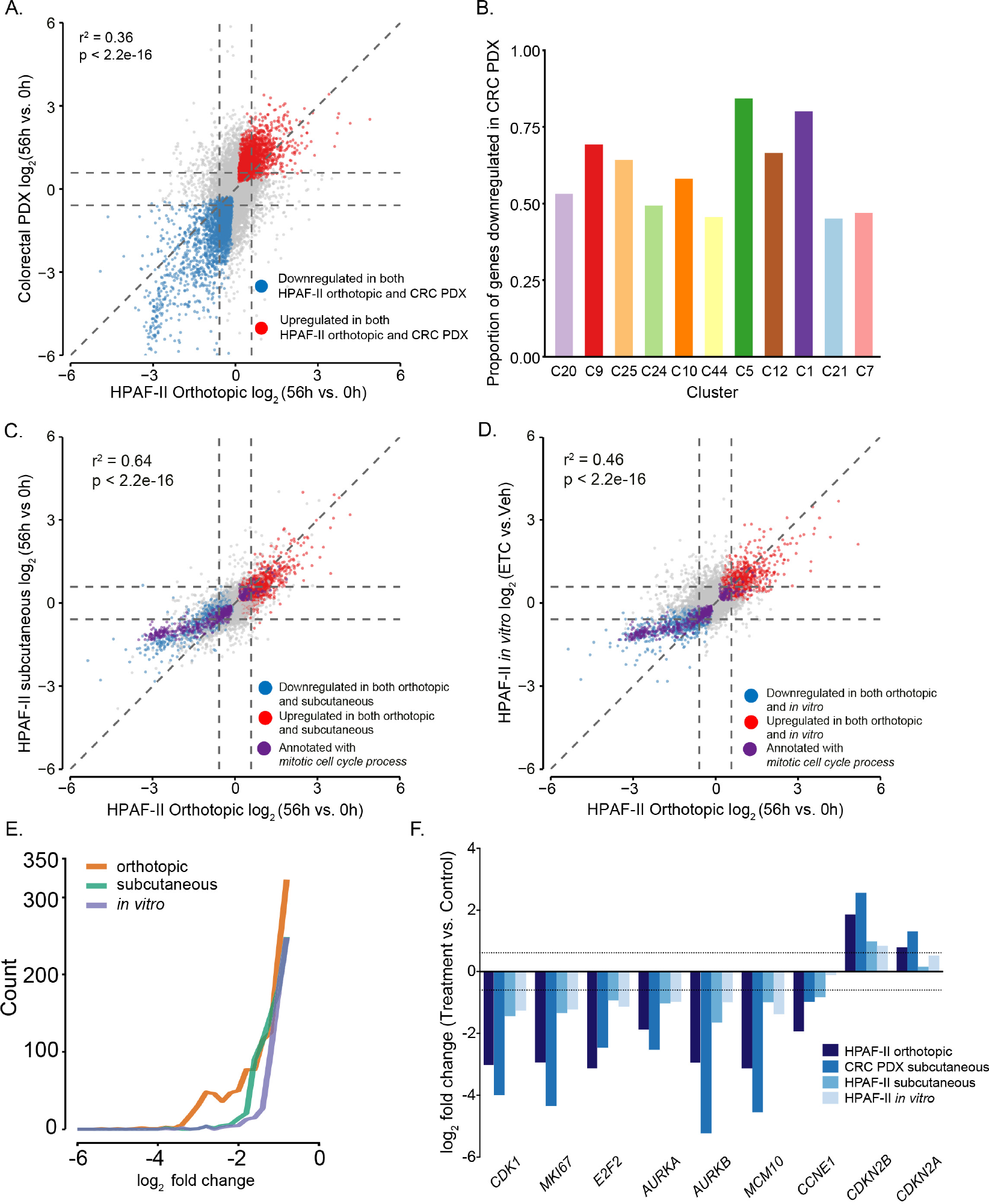
Multi-model comparison identifies core Wnt-activated genes most effectively in pancreatic orthotopic and colorectal PDX models. A. Gene expression changes in the HPAF-II orthotopic xenografts are highly correlated with those in a colorectal patient derived xenograft model (CRC PDX) with a RSPO3 translocation (described in (Madan et al., 2016)) (p < 2.2 × 10^−16^). Diagonal dashed line is where genes would fall if they changed equally in the two models. Horizontal and vertical dashed lines indicate a 1.5 fold change.
B. Genes identified as *Wnt-activated* in the HPAF-II orthotopic model are also significantly downregulated in the CRC PDX. Proportion of genes in each indicated time-series cluster that was also downregulated (FDR < 10%) in the CRC PDX is shown.
C. The HPAF-II orthotopic model is markedly more responsive than a subcutaneous model to Wnt inhibition. Gene expression changes in ETC-159 treated HPAF-II orthotopic compared to subcutaneous xenografts are plotted as in 2A, above. Genes annotated as *mitotic cell cycle processes* (GO BP1903047, purple dots) are far less responsive to PORCN inhibition in the subcutaneous model.
D. The HPAF-II orthotopic model is dramatically more responsive to Wnt inhibition than are HPAF-II cells cultured *in vitro*, data are plotted as in 2C, above. HPAF-II cells in culture were treated for 48 h with 100 nM ETC-159.
E. The distribution of expression changes for the downregulated genes (fold change < 0.5, FDR < 10%) illustrates that the orthotopic model has the most robust response across three different HPAF-II models.
F. Cell cycle gene expression changes were more robust in CRC PDX and HPAF-II orthotopic xenografts compared to subcutaneous and *in vitro* models. Representative cell cycle genes are shown.

The shared response of the Wnt-addicted pancreatic and colorectal cancers led us to evaluate the importance of the stromal microenvironment in regulating the response to Wnt inhibition. We compared the effects of ETC-159 on the transcriptome of HPAF-II cells implanted as orthotopic xenografts, subcutaneous xenografts or cultured *in vitro*. The response to Wnt inhibitor treatment in orthotopic xenografts was correlated to the response in both the subcutaneous model (r^2^ = 0.64, Fig. 2C) and the *in vitro* model (r^2^ = 0.46, Fig. 2D). However, the magnitude of the responses varied widely between the models. Differential expression analysis of the three HPAF-II models identified 4,409 genes whose response to ETC-159 was significantly different (interaction test, FDR < 10%) between models. These genes were enriched for processes including cell cycle, ribosome biogenesis (Fig. S3B). 51% of all *Wnt-activated* genes responded differently to ETC-159 across the three systems, with many more genes showing robust fold changes in the orthotopic model compared with the other systems (Fig. 2E). In particular, cell cycle associated genes changed most robustly in the orthotopic model (Figs 2C-D and F). For example, *AURKA* decreased by 3.7 fold at 56 hours after the start of PORCN inhibition in the orthotopic system, 2.1 fold in the subcutaneous model and only 1.7 fold *in vitro*. *Cyclin E1* did not change *in vitro* but decreased by 1.7 fold and 3.8 fold in the subcutaneous and orthotopic model, respectively (Fig. 2F). Negative regulators of the cell cycle such as *CDKN2B* also exhibited differences in their magnitude of response to PORCN inhibition. In addition to cell cycle associated genes we identified several other genes that did not respond to Wnt inhibition *in vitro* but behaved as WNT targets *in vivo*(i.e. *EPHB3* and *TGFBI*, Fig. S3C). Thus, the overall response to Wnt inhibition was reduced in the subcutaneous model and further blunted *in vitro* (Figs 2C-F).

Further highlighting the importance of the orthotopic model was the considerable difference in baseline gene expression between models ~30% of all expressed genes (4,992 genes) were differentially expressed even before treatment when comparing *in vitro* and orthotopic xenografts. This included greater than two-fold increases (moving from cultured cells to the orthotopic model) in Wnt pathway genes *WNT11*, *WNT2B*, *FZD8*, *FZD4*, and *LRP5*, as well as the Wnt target genes *NKD1*, *SP5*, *LGR5* and *AXIN2* (Table S4). Additionally, 3,515 genes were differentially expressed at baseline when comparing subcutaneous xenografts with the orthotopic model (absolute fold change > 1.5, FDR < 10%) (Figs S3D-E). Not surprisingly, a number of genes more highly expressed in the orthotopic xenografts compared to *in vitro* were associated with multicellularity, innate immunity, extracellular matrix organization and cell adhesion (Fig. S3D).

The much less pronounced effect of ETC-159 on the expression of cell cycle genes in cell culture is consistent with our previous observations that PORCN inhibitors do not inhibit the growth of Wnt-addicted cancer cells in short-term 2D cell culture (Proffitt et al., 2013). This data illustrates that the orthotopic and PDX mouse models more accurately recapitulate the tissue-specific tumor microenvironment and highlights the value of the orthotopic model in identifying core Wnt regulated genes.

### PORCN inhibition leads to early downregulation of MYC and its targets

The time-series clustering analysis (Fig. 1E) identified sets of genes (clusters) having similar dynamics of response to PORCN inhibition, suggesting that each cluster may be regulated by distinct mechanisms. To investigate the differences in the transcriptional regulation of these genes, we performed a Transcription Factor Binding Site (TFBS) motif analysis on the promoters of the *Wnt-activated* genes (Fig. 3A).

**Figure 3:**
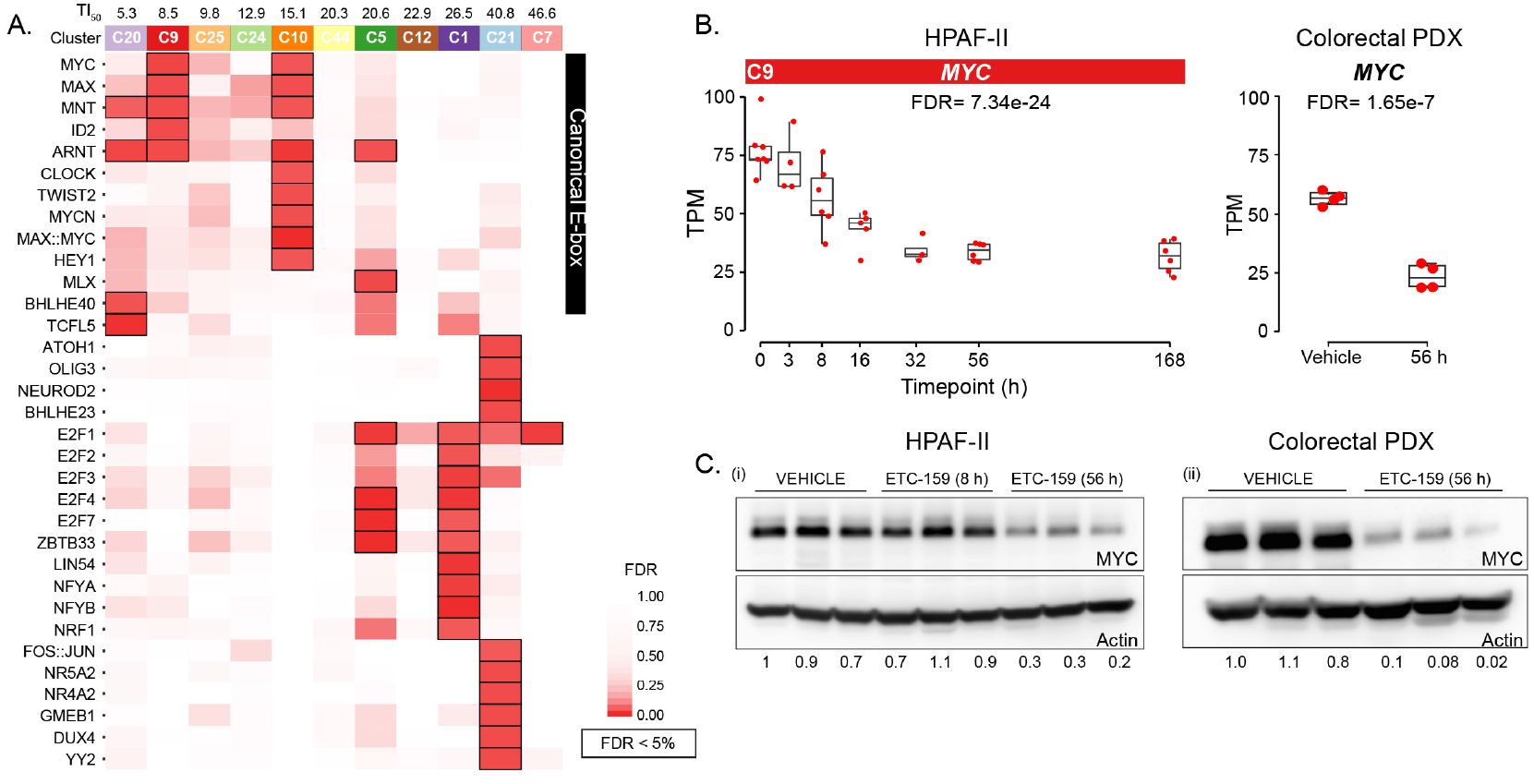
The waves of Wnt-activated genes are associated with distinct sets of transcription factor binding sites with enrichment for E-boxes in the early responding clusters. A. Clusters of *Wnt-activated* genes are enriched (FDR < 5%) for distinct transcription factor binding site motifs. Promoters of genes in each cluster were scanned for motifs present in JASPAR 2016 using using FIMO (Grant et al., 2011), and enrichment for each cluster was calculated. Promoters of genes in the early downregulated clusters are enriched for canonical E-boxes.
B. *MYC* gene expression is partially inhibited by ETC-159 treatment of HPAF-II orthotopic tumors and a colorectal PDX (Madan et al., 2016).
C. MYC protein abundance is reduced in both HPAF-II orthotopic xenografts and colorectal PDX following 56 h treatment with ETC-159. The ratio of MYC protein compared to P-actin protein abundance for each lane is indicated.

Unexpectedly, the promoters of genes downregulated immediately following Wnt withdrawal (e.g. C9, TI_50_ = 8.5 h) did not show significant enrichment for TCF7L2 binding sites (p-value=0.21). The majority of TCF7L2 binding events are found to be intergenic rather than promoter-associated (Fig. S4) (Stevens et al., 2017). The promoters of genes in the most rapidly responding clusters (i.e. C9, C10, C20, C24 and C25, TI_50_ < 20 h) were rather significantly enriched for canonical E-box motifs, bound by transcription factors including MYC, HEY1, CLOCK and ID2 (Fig. 3A). The genes in clusters that responded later (TI_50_ > 20 h) were enriched for E2F, NRF and NFYB binding sites(Dolfini and Mantovani, 2013).

The enrichment of E-box binding motifs occurred in some early responding *Wnt-activated* genes (e.g. C20, C9) whose expression fell even before *MYC* mRNA decreased, suggesting additional levels of regulation (Fig. 3A). Upon PORCN inhibition, *MYC* mRNA responded like a direct WNT target gene with an early and sustained decrease (C9), albeit only to ~50% of its initial mRNA abundance (Fig. 3B). This is consistent with the well-established role of β-catenin signaling in the regulation of *MYC* expression(Myant and Sansom, 2011). In addition to being a transcriptional target of Wnt signaling, MYC protein abundance can be directly regulated by GSK3 by phosphorylating it at Threonine 58, thus priming it for ubiquitylation and proteasomal degradation (Acebron et al., 2014; Arnold et al., 2009; Sears et al., 2000; Taelman et al., 2010). As Wnt signaling inhibits AXIN-associated GSK3, blocking Wnt signaling increases the activity of GSK3 and promotes MYC degradation. Indeed, we observed a more pronounced change in MYC protein (2.8-14.5 fold reduction) than *MYC* mRNA (2.0-2.2 fold reduction) in both HPAF-II orthotopic tumors and the colorectal PDX models 56 h after PORCN inhibitor treatment (Figs 3B-C). These results suggest that the Wnt-dependent decrease in *MYC* transcripts was coupled with post-transcriptional regulation of MYC protein abundance, i.e. a Wnt/STOP effect in Wnt addicted tumors.

### Wnt signaling regulates MYC via WNT/STOP and Wnt/β-catenin pathway

To assess the relative contributions of Wnt-regulated MYC mRNA expression (Wnt/β-catenin) and Wnt/GSK3-regulated MYC protein stability, we generated HPAF-II cell lines stably overexpressing either *Myc* (MYC OE) or GSK3-resistant *Myc* (MYC T58A) under the control of the Wnt-independent CMV promoter. The CMV promoter drove 10-15 fold higher *Myc* mRNA expression in orthotopic tumors (Fig. 4A). We then compared the effect of PORCN inhibition on the growth of HPAF-II, HPAF-II (MYC OE) and HPAF-II (MYC T58A) orthotopic xenografts.

**Figure 4:**
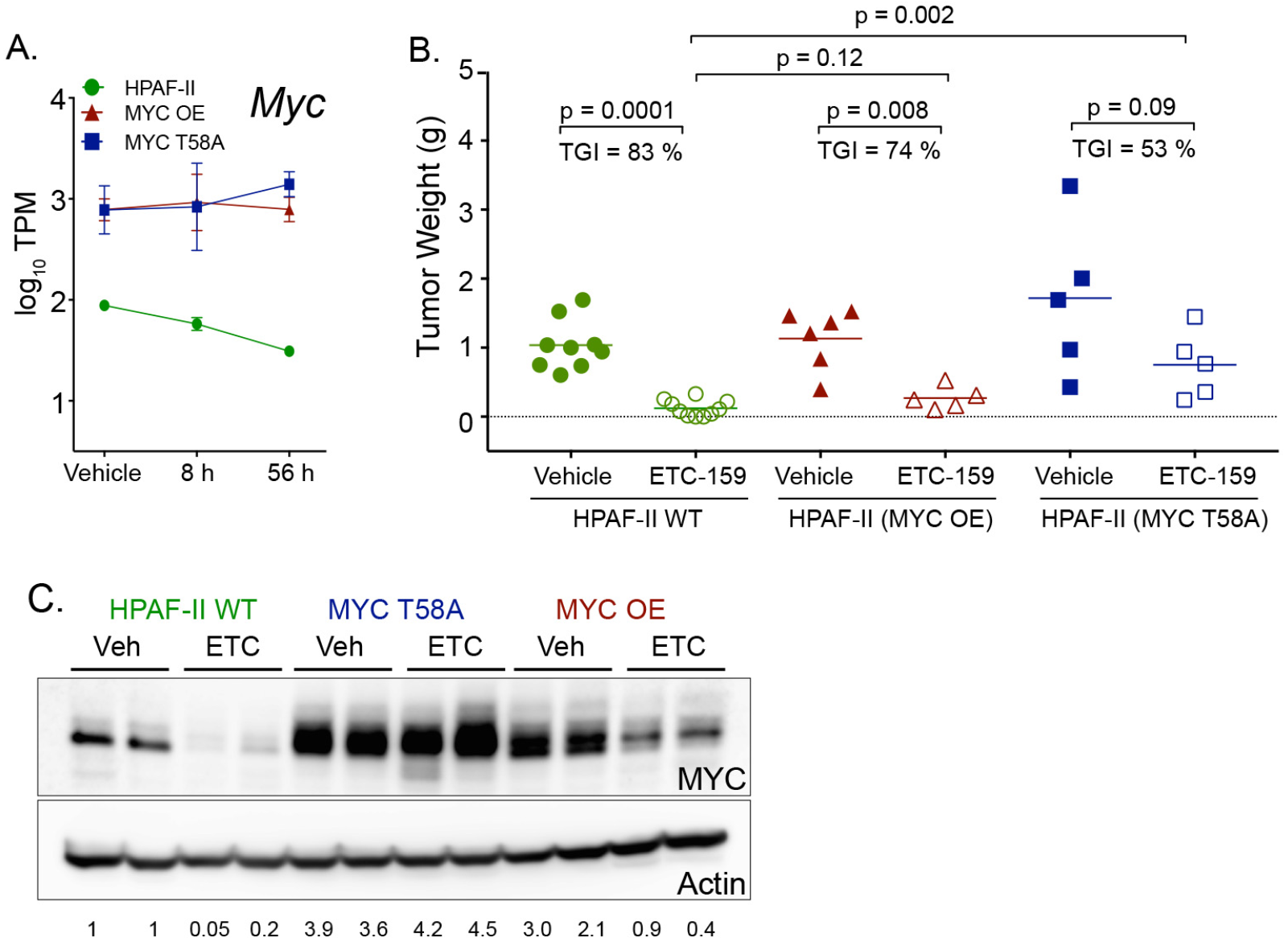
Stabilized MYC partially reverses the effects of Wnt inhibition. A. Expression of non-endogenous *Myc* transcripts in MYC OE or MYC T58A tumors does not change with ETC-159 treatment in contrast to the response of the endogenous *MYC* transcript in the HPAF-II tumors.
B. MYC T58A tumors are partially responsive to Wnt inhibition. HPAF-II cells, MYC OE or MYC T58A HPAF-II cells were injected into the pancreas as before. Following the establishment of tumors, mice were treated daily with 30 mg/kg ETC-159. Tumor weights after 28 days of treatment are shown. N = 5-9 mice/group. Differences between conditions were assessed using Mann-Whitney test (two-tailed). TGI, tumor growth inhibition.
C. Ectopically expressed MYC is sensitive to GSK3-mediated degradation. ETC-159 treatment reduces the protein abundance of both endogenous and ectopically expressed MYC in HPAF-II and MYC OE tumors. Mutation of the GSK3 phosphorylation site prevents the decrease in MYC protein abundance in response to Wnt inhibition in the HPAF-II (T58A) tumors. Ratio of MYC levels compared to β-actin levels for each lane is indicated.

The MYC OE orthotopic tumors had a marked increase in MYC protein but there was no overall increase in tumor growth and remarkably they still responded significantly to PORCN inhibition (Figs 4B-C). Thus, restoration of MYC by overexpression does not rescue tumors from the effects of Wnt inhibition. This could be either because MYC protein is not rate-limiting for tumor growth in this setting, or that PORCN inhibition was able to drive MYC degradation. Consistent with this Wnt/STOP effect, we found that PORCN inhibition caused a decrease in MYC protein abundance despite no change in ectopic *MYC* mRNA levels (Figs 4A, 4C).

We next examined if blocking the Wnt/STOP effect on MYC protein altered the response to PORCN inhibition. The abundance of MYC T58A did not change upon PORCN inhibition (Fig. 4C) and tumors with stabilized MYC grew larger and showed a partial response to PORCN inhibition (Fig. 4B). Taken together, these findings indicate that in addition to its transcriptional regulation, inhibiting Wnt signaling regulates the growth of the tumors by directly regulating MYC protein abundance via a GSK-dependent mechanism. Further, the finding that tumors with stabilized and over-expressed MYC still partially respond to PORCN inhibition shows that Wnts regulate the growth of the pancreatic tumors via both Myc-dependent and Myc-independent pathways. This is similar to findings in the murine intestine, where MYC is essential for the oncogenic effects of APC deletion (Sansom et al., 2007) but alone is insufficient to drive tumorigenesis (Finch et al., 2009).

### Distinguishing between Myc-dependent and Myc-independent regulation of Wnt target genes

To directly identify MYC-independent and MYC-dependent WNT target genes, we performed an additional set of RNA-seq experiments to examine gene expression changes in orthotopic tumors generated from HPAF-II WT, MYC OE or MYC T58A cells. We selected as the time points 0, 8 and 56 hours after the start of therapy to allow us to examine early direct targets of both Wnt/β-catenin and Wnt/GSK3/MYC signaling. Consistent with the previous experiment, ETC-159 treatment reduced MYC protein by ~70% in 56 h in HPAF-II WT tumors, while protein levels in MYC T58A tumors showed no decrease (Fig. 5A).

**Figure 5:**
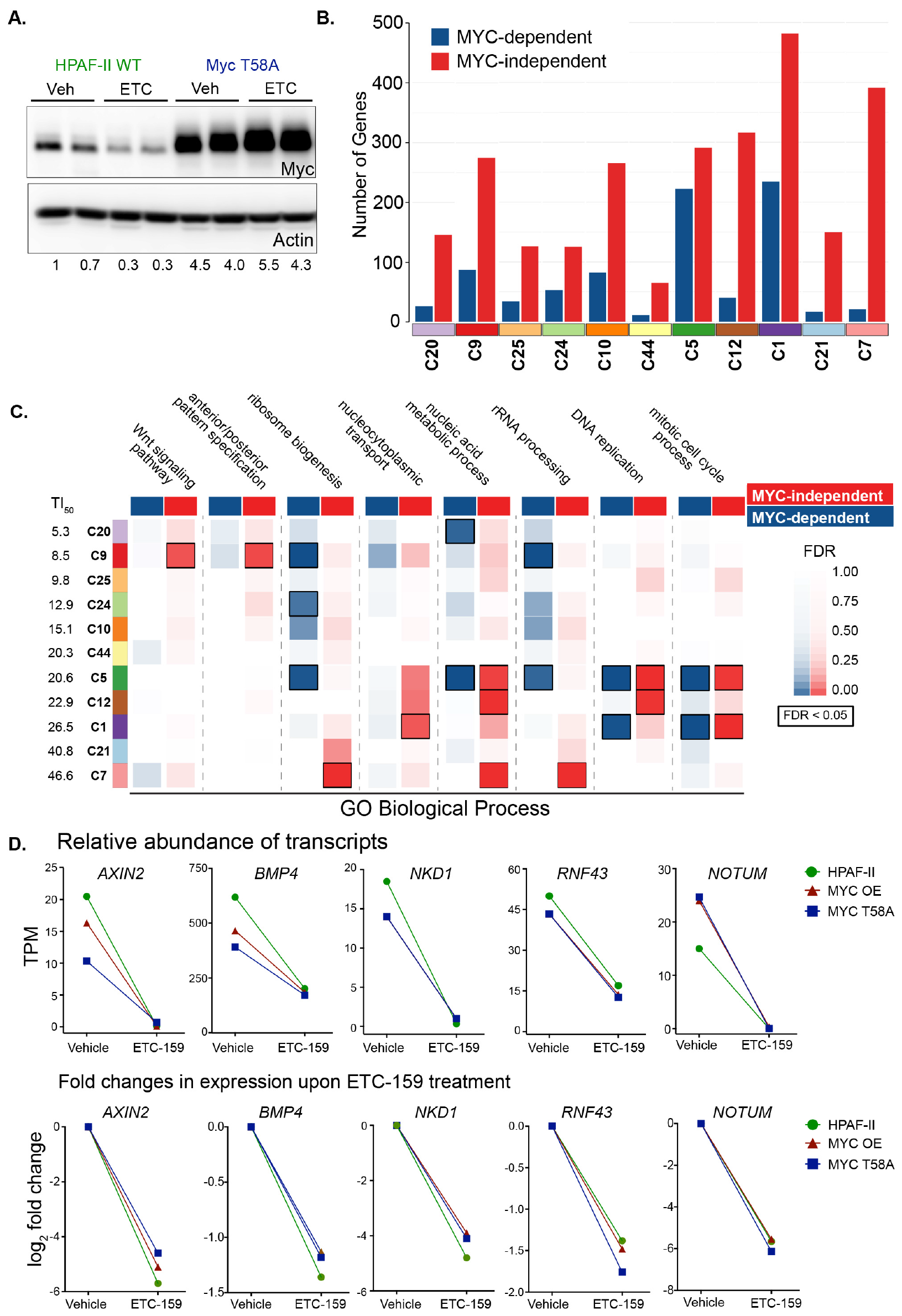
The majority of early responding Wnt-activated genes are Myc-independent. A. Treatment with ETC-159 for 56 h reduces the protein abundance of MYC in HPAF-II WT xenografts but not HPAF-II T58A xenografts. Ratio of MYC levels compared to P-actin levels for each lane is indicated.
B. The majority of *Wnt-activated* genes are Myc-independent. *Wnt-activated* genes were classified as either Myc-dependent or Myc-independent based on whether they responded differently to ETC-159 treatment (interaction test, q-value < 10%) across the three xenograft models studied (HPAF-II, MYC OE and MYC T58A).
C. MYC-dependent and -independent *Wnt-activated* genes in each time-series cluster regulate distinct biological processes (GO:BP). Annotated WNT target genes (i.e. Wnt signaling and anterior/posterior patterning) are MYC-independent. Ribosomal biogenesis and cell cycle genes are regulated both by MYC-dependent and -independent pathways.
D. Representative examples of early responding MYC-independent *Wnt-activated* target genes. Expression and log_2_ fold changes in HPAF-II, MYC OE and MYC T58A xenograft model systems treated with ETC-159 for 56 hours.

We identified 2,131 genes whose transcriptional response to PORCN inhibition *in vivo* was dependent on MYC status (FDR < 10%, Table S5). These genes, whose response to PORCN inhibition was different between WT, MYC OE or MYC T58A tumors, were classified as MYC-dependent Wnt target genes. Of these genes, 827 (23%) were found in our set of *Wnt-activated genes*. In each of the clusters of *Wnt-activated* genes we determined the fraction that exhibited MYC-dependent or MYC-independent responses (Fig. 5B). The majority of genes that were downregulated most rapidly upon Wnt inhibition (i.e C9, C10, C20, and C25, TI_50_ < 20 h) (Fig. 1E) were MYC-independent. Selected examples of well-established Wnt-regulated MYC-independent genes such as *AXIN2* and *NKD1* are illustrated in Fig. 5D, examining both the relative transcript abundance (top panel) and the log fold changes (bottom panel). Not surprisingly, the MYC-independent Wnt target genes in C9 were associated with Wnt signaling pathways and embryonic patterning (Fig. 5C).

Only third wave clusters C5 and C1, changing with TI_50_ of 20.6 and 26.5 h, contained a sizeable fraction (> 25%) of MYC-dependent genes (Fig. 5B) that were enriched in ribosome biogenesis and cell cycle processes discussed below. We did not observe enrichment for E-boxes in the clusters C1 and C5 (Fig. 3A). Interestingly, the subset of genes in these clusters that were MYC-dependent were also not specifically enriched for MYC TFBS, suggesting either they are indirect targets of MYC or that TFBS analysis is not powerful enough to detect a clear enrichment for these MYC motifs.

### Regulation of cell cycle by Wnts in Wnt-addicted cancers is Wnt/GSK3 dependent

Our initial analysis (Fig. 1) demonstrated that cell cycle and ribosomal biogenesis are two key pathways that are transcriptionally regulated by Wnt inhibition, with multiple genes regulating cell cycle changing in a time-dependent manner, including *CDK1*, *E2F2*, *E2F1*, *CDKN2B* and *CDKN2A* (Fig. 6A). Consistent with this robust regulation of cell cycle genes, Ki67 positive cells were significantly reduced in the tumors as early as 56 hours after starting ETC-159 and were further reduced at 7 days of treatment (Fig. 6B).

**Figure 6:**
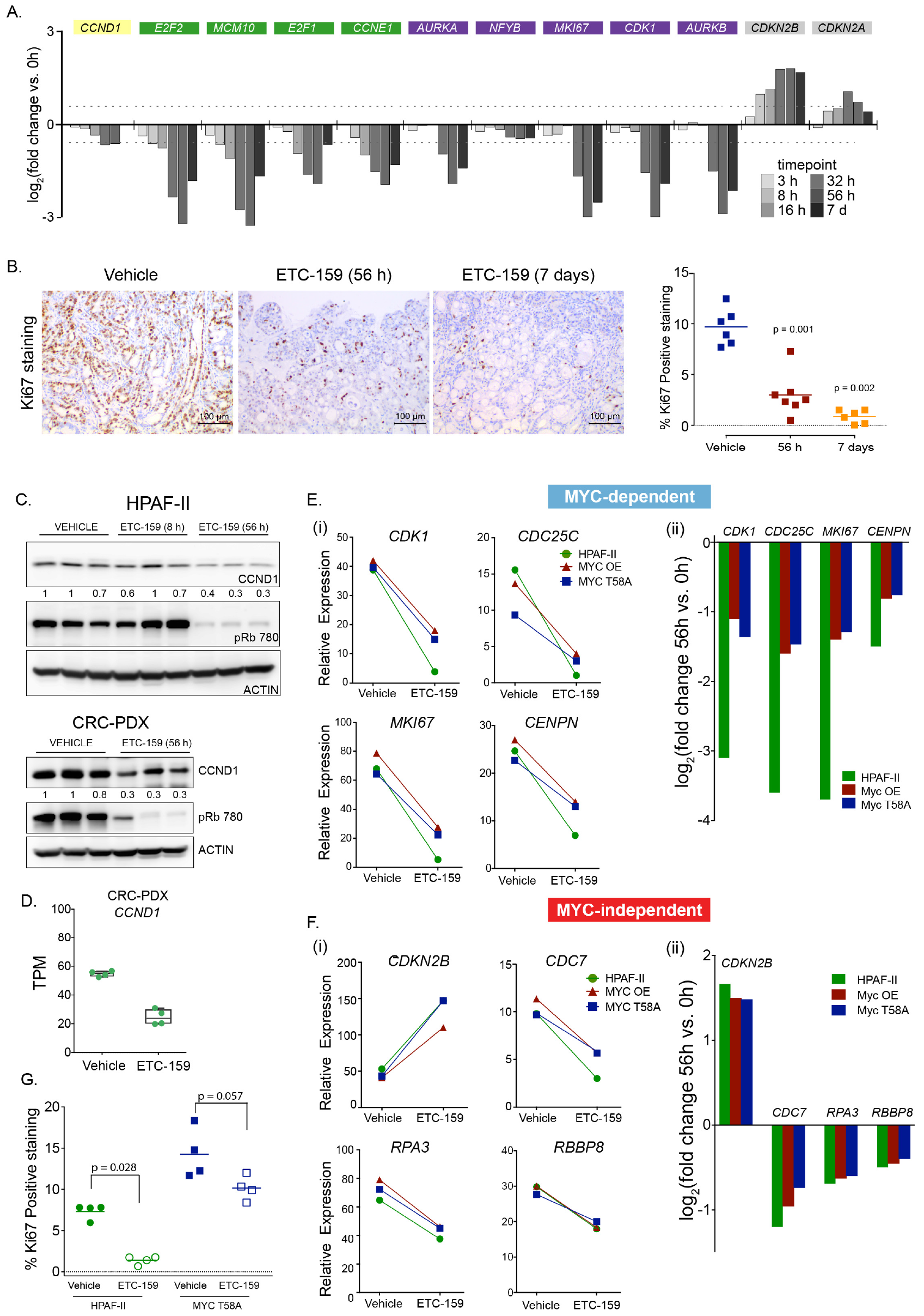
Wnt regulated cell cycle changes are only partially influenced by Myc. A. Robust changes in the expression of representative cell cycle genes over 7 days of PORCN inhibition in HPAF-II orthotopic xenografts. The dotted line is at 1.5 fold (log_2_ 0.58).
B. Significant decrease in proliferating cells over time. Ki67 positive cells (left) were quantitated (right) on an entire section that was scanned and analyzed using NIS-Elements software.
C. CCND1 protein changes more robustly compared to mRNA. *CCND1* mRNA decreases 33% (see 6A, and 6D) while CCND1 protein decreases ~65% after 56 h treatment in both HPAF-II and CRC PDX models. Each lane is from an independent tumor. Ratio of CCND1 levels compared to the β-actin levels for each lane is indicated. ETC-159 treatment reduces levels of pRb (S780) in the both HPAF-II tumors and Colorectal PDX.
D. Changes in the expression of CCND1 after 56h in CRC PDX model.
E. Representative cell cycle genes whose expression is influenced by MYC overexpression (i) plotted as relative expression (TPM), (ii) plotted as log_2_ fold change.
F. Representative cell cycle genes whose expression is independent of MYC overexpression. (i) plotted as relative expression (TPM), (ii) plotted as log_2_ fold change.
G. MYC stabilization only partially rescues proliferation upon PORCN inhibition. Ki-67 staining quantitated as in B above.

Genes associated with mitotic cell cycle processes and DNA replication were enriched in the third wave of clusters (C1, C5, C12; TI_50_ 20.6-26.5 hrs) (Fig. 1F). These clusters were enriched for binding sites for the E2F and NFY families of transcription factors (Fig. 3A) that cooperatively regulate cell cycle genes (Dolfini and Mantovani, 2013; Ly et al., 2013; van den Heuvel and Dyson, 2008).

Interestingly, *E2F1* and *E2F2* gene expression decreased at the same rate as the other cell cycle genes, suggesting that the early decrease in the expression of cell-cycle related genes was not due to changes in these *E2F* mRNAs (Fig. 6A). E2F activity is also regulated by cyclin dependent kinase signaling through p105/Rb. Indeed, the expression of the CDK inhibitors increased as early as 8 h after PORCN inhibition in orthotopic tumors, associated with a subsequent decrease in Rb phosphorylation (Figs 6A and C). Thus, CDK inhibition and decreased Rb phosphorylation is likely to be a major mechanism driving the decrease in the transcription of E2F target cell cycle genes.

Notably, the abundance of cell cycle regulators such as Cyclin D1 and E1, amongst others, is also regulated by Wnt/STOP signaling (Acebron et al., 2014). Similar to MYC, after 56 h of ETC-159 treatment the protein abundance of CCND1 was reduced by ~3 fold (Fig. 6C) while *CCND1* transcript levels were only reduced by ~1.5 fold in both HPAF-II xenografts and colorectal PDX (Figs 6A and D). Thus, our *in vivo* data in Wnt driven cancers support the data from *in vitro* studies (Acebron et al., 2014) that Wnt signaling regulates CCND1 and MYC by both transcriptional and post-transcriptional mechanisms.

We further examined the role of MYC in the regulation of cell cycle genes. Notably, MYC overexpression had no effect on baseline expression of the cell cycle genes and Wnt inhibition reduced their expression, albeit to differing extents even in the presence of stabilized MYC (Figs 6E-F). Stabilized MYC blunted the effect of PORCN inhibition on the expression of a subset of the cell cycle genes, e.g. *CDK1*and *MKI67* (Fig. 6E). However, a number of other cell cycle genes (e.g. *CDKN2B*, *CDC7*, *RBBP8* and *RPA3*) were MYC-independent and responded to Wnt inhibition even in MYC-stabilized tumors (Fig. 6F). Consistent with the observed transcriptional response, there was a partial reduction of Ki67 staining in ETC-159 treated MYC-stabilized tumors (Fig. 6G). Taken together these findings indicate that Wnt regulates the cell cycle in cancers via multiple pathways, both dependently and independently of MYC, and through both transcriptional and Wnt/STOP mechanisms.

### Wnt signaling regulates Ribosomal Biogenesis

The enrichment for rRNA processing and ribosomal biogenesis in the Wnt activated gene clusters (Fig. 1F) suggested that PORCN inhibition would lead to a reduction in ribosome formation and protein synthesis. Indeed, nearly all genes encoding ribosomal protein subunits (*RPSs* and *RPLs*) were downregulated (Fig. 7A), with 94% of differentially regulated *RPSs* and *RPLs* being present in late responding clusters, C7 (TI_50_ 46.6 h) or C1 (TI_50_ 26.5 h). Although the expression of *RPS* and *RPL* genes was reduced by only ~30-40 %, the changes were largely coherent albeit with some outliers of unknown significance. The changes were more apparent following 32 h of treatment, suggesting that these genes are indirectly regulated by Wnt signaling. We next examined if this was reflected in the abundance of ribosomal subunit proteins. In a parallel mass spectrometry experiment that only detected high abundance proteins (see Methods), we confirmed that the RPS and RPL proteins were also coherently downregulated at 56 h (Fig. 7B, Table S6). Given the high abundance of ribosomal proteins, this suggests a dramatic shift in ribosome biogenesis.

**Figure 7:**
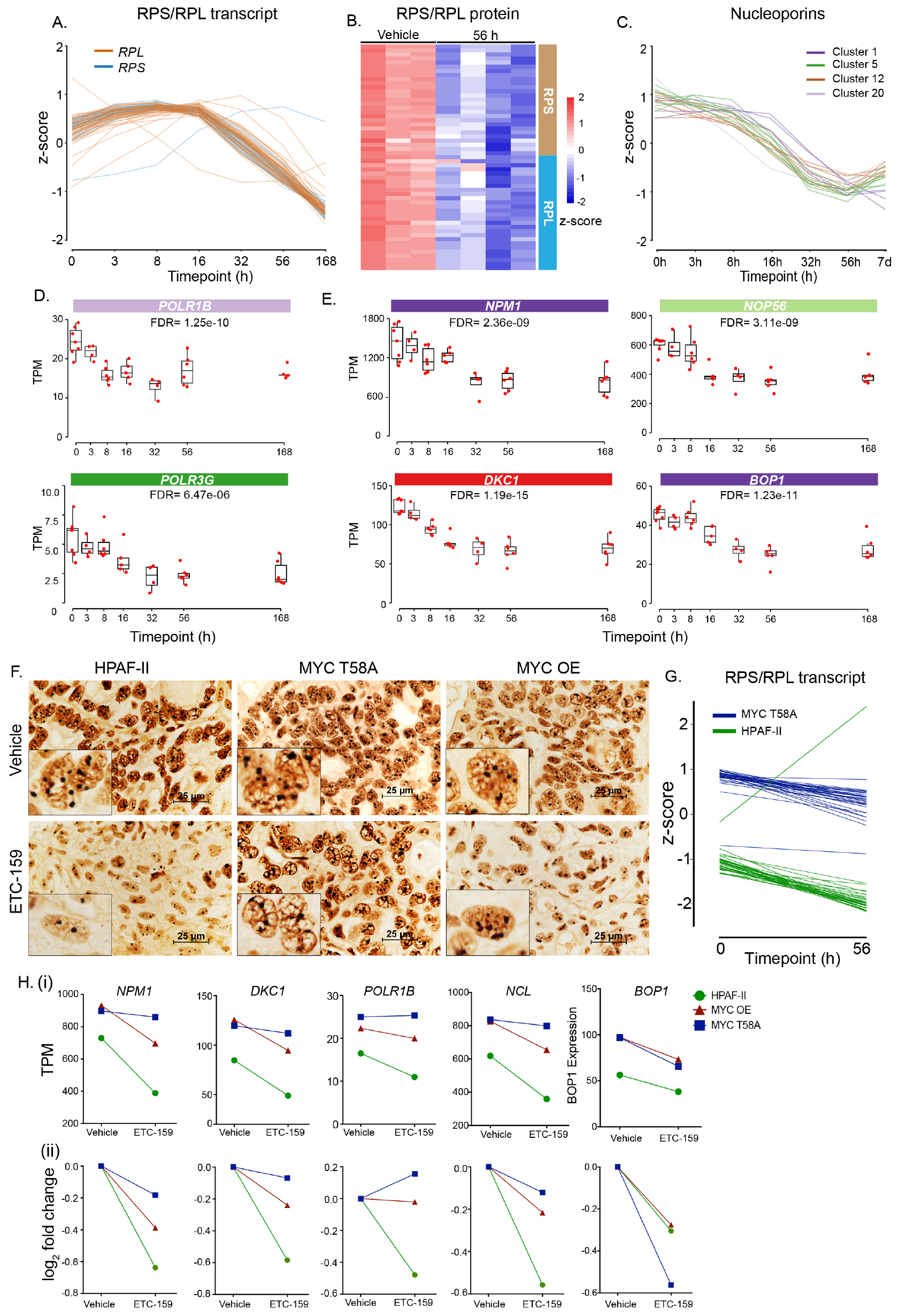
Wnt and Myc co-regulate ribosome biogenesis. A. Coherent changes in the expression of genes encoding ribosomal proteins (*RPLs*/*RPSs*) over time following PORCN inhibition.
B. Coherent changes in the abundance of ribosomal proteins over time as assessed by mass spectrometry following PORCN inhibition.
C. Coherent changes in the expression of nucleoporin genes over time following PORCN inhibition.
D. Gene expression of RNA polymerases subunits, including *POLR1B* and *POLR3G*, that transcribe ribosomal RNA are Wnt regulated.
E. Key regulators of ribosome biogenesis *NPM1, BOP1, NUP58* and *DKC1* change over time following PORCN inhibition.
F. Wnt inhibition reduces nucleoli but this is rescued by expression of stabilized MYC. Changes in nucleolar size and abundance in tumor sections was assessed by silver staining.
G. Expression of ribosomal subunit genes (RPLs/RPSs) is enhanced by stabilized MYC T58A but remains sensitive to PORCN inhibition. The outlier in HPAF-II tumors is RPS27L, as in 7A.
H. Representative *Wnt-activated* ribosome biogenesis genes that are MYC-dependent. Changes in selected genes in MYC OE and T58A tumors are shown. (i) plotted as relative expression (TPM), (ii) plotted as log_2_ fold change.

Ribosomal biosynthesis requires multiple processes including nucleocytoplasmic transport and rRNA expression and processing (van Riggelen et al., 2010). We found that genes required for nucleocytoplasmic export including exportins and nucleoporins were similarly coherently downregulated implicating the regulation of ribosome assembly by Wnt signaling (Fig. 7C). Multiple components of the machinery required for rRNA transcription, including several subunits of RNA polymerases *POLR1* and *POLR3* (Fig. 7D) and rRNA processing factors (e.g. *NPM1*, *DKC1*) were also downregulated (Fig. 7E). Finally, consistent with the changes in gene and protein expression, the size of nucleolar organizer regions was reduced by ETC-159 treatment (Fig. 7F). Taken together, these data indicate that ribosomal biogenesis is globally regulated by Wnt signaling. A global decrease in protein synthesis coupled with a halt in the cell cycle likely explains how PORCN inhibition blocks tumor progression in Wnt-addicted cancers (Ruggero and Pandolfi, 2003).

We asked if Wnt regulation of ribosome biogenesis was explained by its effect on MYC, a recognized regulator of ribosome biogenesis (van Riggelen et al., 2010). In contrast to the cell cycle genes, the baseline expression of ribosome subunit and biogenesis genes was increased by stabilized MYC (Figs 7G-H). However, a number of these genes remained sensitive to PORCN inhibition and decreased after 56 h of ETC-159 treatment even in cells with MYC T58A (Fig. 7G). These Wnt-regulated, Myc-independent ribosome genes includes virtually all of the *RPLs* and *RPSs* (C1 and C7) (Figs. 7G and S5). Another subset of genes involved in rRNA synthesis and processing (e.g., *NPM1*, *DKC1*, *POLR1B*) were MYC-dependent WNT target genes. These genes were both highly MYC responsive at baseline, and consistent with their regulation by Wnt/STOP regulation of MYC protein abundance, did not respond to Wnt inhibition if MYC T58A was present (Fig. 7H). These MYC-dependent genes are enriched for E-boxes in their promoters.

Our analysis thus establishes a key role of Wnt signaling in ribosome biogenesis via two routes. One route, via MYC, is regulated both through Wnt-driven *MYC* expression and via the Wnt/STOP pathway. The other route is MYC-independent and is a downstream effect of WNT signaling on the transcription of ribosomal genes.

## Discussion

The development of targeted drugs that rapidly and robustly inhibit PORCN provides a unique opportunity to examine in real time the consequences of Wnt withdrawal in Wnt-addicted human cancers. This time-based analysis of Wnt signaling and its interaction with MYC, provides a comprehensive assessment of the role played by Wnt ligands in driving Wnt-addicted cancer. Importantly, the high concordance of the transcriptional changes in Wnt-addicted RSPO3-mutant colorectal and RNF43-mutant pancreatic cancers reveals core shared pathways regulated by Wnt signaling in cancer. Previous studies examining the targets of Wnt signaling in cancer have focused on models that are driven by loss of function mutations in APC. Here, the use of Wnt ligand driven cancer mouse models casts a broader net, identifying a unexpectedly large number of genes whose expression depends on continued presence of Wnt ligand many of which are independent of β-catenin.

The genes whose expression changes most rapidly after PORCN inhibition, the early wave clusters, were predictably enriched for well-established β-catenin target genes (Moon and Gough, 2016). However, our analysis revealed a large number of co-regulated genes that were not known β-catenin targets. DNA sequence-based analysis of enrichment for TCF/LEF binding sites was not a useful approach to discriminate if these early changing genes could be additional β-catenin targets or they could be regulated by multiple non-canonical pathways. Indeed, while many individual studies find TCF/LEF sites in the promoters of selected genes, our findings support the results from genome-wide analyses showing that functional TCF/LEF sites are often present at large distances from transcriptional start sites (Ramakrishnan and Cadigan, 2017). Additionally, recent studies have established that even β-catenin promoter binding is not sufficient to identify β-catenin transcriptionally regulated genes^5, 40^.

Interestingly our analysis revealed that E-box transcription factor binding sites are enriched in the early changing genes, followed at later time points by enrichment for E2F-binding sites. Finally, the fourth wave of genes was enriched for a broader set of TFBS that are likely to be regulated as secondary, downstream events. The enrichment for E-boxes strongly suggested a role for MYC. *MYC* is a potent oncogene and its activation is a hallmark of cancer initiation and maintenance (Dang, 2010; Gabay et al., 2014). MYC is required for tumorigenesis following β-catenin activation by APC loss in the gut but not in the liver (Reed et al., 2008; Sansom et al., 2007). Hence, it was an open question if MYC would be important downstream of *RNF43* mutations in pancreatic cancers, where many additional pathways are activated by the Wnt addiction (Sansom et al., 2007; Wilkins and Sansom, 2008). Using a model of Wnt-addicted human cancer with stabilized MYC we were able to disentangle the interaction of Wnt and MYC and stratify the role of Wnts and MYC in regulating cell cycle and ribosomal biogenesis. One notable difference was that stabilization of MYC did not enhance the expression of cell cycle genes. However, stabilized MYC could partially overcome the effect of Wnt inhibition on expression of a subset of cell cycle genes (Annibali et al., 2014). Whereas, *Myc* overexpression and stabilization more profoundly affected genes regulating various processes associated with ribosomal biogenesis. Here too, the response to Wnt inhibition was variable as a large subset of genes, including ribosomal proteins, responded to PORCN inhibitors with similar fold changes, while others were “immune” to Wnt inhibition in the presence of stabilized MYC. This suggests a complex interaction of MYC and Wnt-regulated pathways driving these processes.

Ribosomes are overexpressed in cancer and have become novel targets for anticancer therapies, for instance, by triggering nucleolar stress (Pelletier et al., 2018; Quin et al., 2014; Ruggero and Pandolfi, 2003; Sulima et al., 2017). While MYC is known to regulate ribosome biogenesis (van Riggelen et al., 2010), the role of Wnts has been less clear (Kraushar et al., 2015; Pfister and Kühl, 2018). Here we show for the first time that Wnt signaling globally impacts multiple steps in ribosomal biogenesis both directly and by regulating the transcription and protein abundance of Myc via the Wnt/STOP pathway and this is shared in both Wnt-addicted pancreatic and colorectal cancers.

Comparing the effect of PORCN inhibition across different models confirmed the value of studying Wnt signaling in an orthotopic microenvironment or in the present of native stroma (CRC PDX). The experimental value of the orthotopic model using a cell line is that it is more amenable to genetic manipulation such as the introduction of stabilized MYC, allowing a more detailed analysis of the role of downstream drivers.

The stabilization of MYC via the Wnt/GSK3 signaling axis highlights how this mechanism can target MYC and other cell cycle proteins in cancer, impacting aberrant cell growth (Dang et al., 2017). The Wnt/STOP pathway is likely to have multiple additional targets (Koch et al., 2015; Taelman et al., 2010) that may also play a role in these pancreatic and colorectal cancers. Future studies with high-resolution mass spectrometry at early time points after Wnt inhibition may facilitate their identification. The data provided in this study can facilitate biomarker discovery for patients suffering from Wnt addicted cancers and provides a significant resource for the Wnt and cancer community.

## Material and Methods

### Tumor growth and mice treatment

Mouse xenograft models from HPAF-II cells were established by orthotopic injection of HPAF-II cells in NOD scid gamma mice as described in the supplemental methods.

### Western Blot analysis

Tumors were homogenized in 4% SDS buffer and proteins were resolved on 10% SDS-polyacrylamide gel. Western blots were performed according to standard methods.

### Immunohistochemistry and AgNOR staining

Formalin fixed and paraffin embedded tissue sections were then stained with hematoxylin and eosin, Ki67 or nucleolar organizer regions using standard protocol. Images were acquired using Nikon E microscope.

### RNA isolation and Data analysis

Tumors were homogenized in RLT buffer and total RNA was isolated using RNAeasy kit (Qiagen) according to manufacturer’s protocol. The RNA-seq libraries were prepared using the Illumina TruSeq stranded Total RNA protocol with subsequent PolyA enrichment. Details for QC and data processing for RNA-seq, transcription factor binding site analysis, time series clustering and ChIP-seq analysis are provided in the supplemental methods.

### Proteomics

Tumors were homogenized on dry ice and solubilized with 8M urea and 20 mM HEPES. Trypsin digested Peptides in 0.1% TFA were separated using an Ultimate 3000 RSLC nano liquid chromatography system coupled to a Q-Exactive mass spectrometer. Following data-dependent acquisition, raw data files were loaded and analyzed using Progenesis QI.

## Acknowledgements

We acknowledge the assistance of Yunka Wong and other members of the Virshup lab and members of Experimental Therapeutics Centre. We acknowledge Ralph Bunte, D.V.M., for his expert advice with histological analysis, and the assistance of the vivarium staff including Hock Lee.

This research is supported in part by the National Research Foundation Singapore and administered by the Singapore Ministry of Health’s National Medical Research Council under the STAR Award Program to D.M.V. E.P. acknowledges the support of the MRC London Institute of Medical Sciences, Imperial College, London.

## Author Contributions

Babita Madan, Nathan Harmston, Enrico Petretto and David M. Virshup designed the study. Babita Madan and Gahyathiri Nallan performed the animal studies and biochemical analysis. Nathan Harmston designed and performed the bioinformatics analysis. Alex Montoya and Peter Faull performed the mass spectrometry. Enrico Petretto and David M. Virshup supervised the study. Babita Madan, Nathan Harmston, Enrico Petretto and David M. Virshup wrote the manuscript.

## Supplementary Information

### Material and Methods

#### Animal care

NOD scid gamma mice were purchased from InVivos, Singapore or Jackson Laboratories, Bar Harbor, Maine. The Duke-NUS Institutional Animal Care and Use Committee approved all the animal studies, which complied with applicable regulations. Animals were housed in standard cages and were allowed access *ad libitum* to food and water.

#### Tumor growth and mice treatment

HPAF-II cells were obtained from ATCC. Mouse xenograft models were established by orthotopic injection of HPAF-II cells in NOD scid gamma mice. 1×10^6^ HPAF-II cells stably expressing luciferase, resuspended in 50% matrigel were used for orthotopic injection into the pancreas of NOD-scid-gamma (NSG) mice. The growth of the orthotopic tumors was monitored weekly by injecting the mice with a 150 mg/kg D-Luciferin (Caliper # 122796) and imaging mice using an IVIS imaging system (Caliper Lifesciences). Mice were treated with ETC-159 after establishment of tumors. ETC-159 was formulated in 50% PEG 400 (vol/vol) in water and administered by oral gavage at a dosing volume of 10 μL/g body weight. At sacrifice, tumors were resected, weighed and snap frozen in liquid nitrogen or fixed in 10% neutral buffered formalin. MSCV Myc IRES GFP plasmid was a gift from Scott Lowe (Addgene plasmid # 18770). The T58A site in MSCV Myc IRES GFP plasmid was mutated using site directed mutagenesis. These plasmids were used for generating HPAF-II cells stably overexpressing MYC or MYC T58A under the control of CMV promoter.

#### Western Blot analysis

Tumors were homogenized in 4% SDS buffer using a polytron homogenizer. Equal amount of proteins were resolved on 10% SDS-polyacrylamide gel and transferred to PVDF membranes. Western blots were performed according to standard methods. Myc antibody (cat# ab32072) was obtained from Abcam and anti-rabbit IgG-HRP (cat# P0448) antibody was obtained from Cell Signaling Technology (Danvers, MA). The blots were developed using SuperSignal West femto substrate (Thermo Scientific; Rockford, IL). The images were captured digitally using the LAS-3000 Life Science Imager (Fujifilm; Tokyo, Japan). For stripping and reprobing the blots “Restore western blot stripping buffer” from Thermo Fisher was used.

#### Immunohistochemistry and AgNOR staining

Tumors were fixed in 10% neutral buffered formalin and embedded in paraffin. Tissue sections were deparaffinized in xylene and rehydrated in a series of ethanol gradients. The sections were then stained with hematoxylin and eosin using standard protocol. For Ki67 staining, after antigen retrieval with sodium citrate buffer pH 6.0 for 30 min, the endogenous peroxidase activity was blocked by incubation with H_2_O_2_. The sections were then incubated overnight with Ki67 antibody (Leica Novocastra, Cat #: NCL-Ki67-MM1) followed by incubation with an HRP conjugated secondary antibody for 1 hour. Incubation with 3,3’-diaminobenzidine chromogen substrate resulted in brown staining of Ki67 positive cells and the nuclei were counterstained with Mayer’s hematoxylin. Brightfield images were acquired on a Nikon Eclipse Ni-E microscope. Total percentage of Ki67 positive cells in each sample were quantified using the analysis software from Nikon.

For staining the nucleolar organizer regions, deparaffinized tissue sections were reduced using freshly prepared 1% dithiothreitol solution for 15 minutes. The tissue sections were incubated for 25 minutes in dark with 50% silver nitrate solution prepared in 2% gelatin and 1% formic acid. After staining, the slides were thoroughly washed with distilled water and incubated in 5% sodium thiosulfate solution for 5 min followed by several washes in water. The sections were dehydrated and mounted using DPX (Trerè, 2000). Images were acquired using Nikon E microscope.

#### RNA isolation

Tumors were homogenized in RLT buffer using a polytron homogenizer and total RNA was isolated using RNAeasy kit (Qiagen) according to manufacturer’s protocol. The RNA-seq libraries were prepared using the Illumina TruSeq stranded Total RNA protocol with subsequent PolyA enrichment.

#### RNA-seq analysis

##### Data processing and QC

Sequences were assessed for quality and reads originating from mouse (mm10) were removed using Xenome (Conway et al., 2012). The remaining reads were aligned against hg38 (Ensembl version 79) using STAR v2.5.2 (Dobin et al., 2013) and quantified using RSEM v1.2.31 (Li and Dewey, 2011). Reads mapping to chrM or annotated as rRNA, snoRNA and snRNA were removed. Genes which had less than 10 reads mapping on average over all samples were removed. We also required that genes had more than 10 reads mapping to them in at least 2 replicates across at least 3 conditions. Differential expression analysis was performed using DESeq2 (Love et al., 2014). Independent filtering was not used in this analysis. Pairwise comparisons were performed using a Wald test. To control for false positives due to multiple comparisons in the genome-wide differential expression analysis, we used the false discovery rate (FDR) that was computed using the Benjamini-Hochberg procedure. Previously published CRC PDX RNA-seq data (Madan et al., 2016) (GSE69687) were re-analyzed using the same processing pipeline (above). Expression levels of genes were calculated by converting to counts to TPM (Transcripts per Million).

##### Comparison of transcriptional response across different models

The transcriptional response to PORCN inhibition across the different models was assessed by investigating the expression of protein-coding genes (after removing histone genes) and their response across three different datasets; orthotopic (0 h and 56 h), subcutaneous (0 h and 56 h) and *in vitro* (48 h ETC and 48 h Veh). Reads originating from mouse (mm10) were removed using Xenome for both the orthotopic and subcutaneous dataset. The reads originating from human aligned against hg38 (Ensembl version 79) using STAR v2.5.2 and quantified using RSEM v1.2.31. Reads mapping to chrM were removed. Genes which had less than 10 reads mapping on average over all samples were removed. DESeq2 was used to identify genes that responded differently to treatment depending on the model, i.e. we performed an interaction test in which the the response is modeled as follows: y ~ model + condition + <u>model:condition</u>, where model is orthotopic, subcutaneous or *in vivo* and condition is untreated or treated with ETC-159. Pairwise comparisons to examine baseline differences between models were performed using a Wald test. The false discovery rate (FDR) was computed using the Benjamini-Hochberg procedure.

##### Identification of Myc-dependent and Myc-independent genes

Different models were compared by investigating the expression genes and their response across time in the three different conditions; WT, MYC OE and MYC T58A. Reads originating from mouse (mm10) were removed using Xenome for both the orthotopic and subcutaneous dataset. The reads originating from human aligned against hg38 (Ensembl version 79) using STAR v2.5.2 and quantified using RSEM v1.2.31. Reads mapping to chrM or annotated as rRNA, snoRNA and snRNA were removed. Genes which had less than 10 reads mapping on average over all samples were removed. DESeq2 was used to to identify genes that responded differently to treatment depending on MYC status, i.e. we performed an interaction test in which the response is modeled as follows: y ~ model + timepoint + <u>model:timepoint</u>, where model is WT, MYC OE or MYC T58A and timepoint is 0h, 8 h, 56 h. Pairwise comparisons to examine baseline differences between models were performed using a Wald test. P-values were corrected for multiple testing using the q-value method (Storey and Tibshirani, 2003), with genes with an interaction q-value < 0.1 being classified as MYC-dependent.

#### Time series clustering

Gene-level counts were transformed using a variance-stabilising transformation and converted to z-scores. Time was transformed using a square root transformation. All genes differentially expressed over time (DESeq2, false discovery rate (FDR) < 10%) were clustered using GPClust (Hensman et al., 2015) using the Matern32 kernel with a concentration (alpha) parameter of 0.001 and a length scale of 6.5. Genes were assigned to a cluster which they had the highest probability of being a member of. Several parameter settings were used to assess the stability and robustness of clusters generated, with GPClust found to generate stable clustering across different thresholds and runs. Clustering was performed 100 times for a specified set of parameters, with the best clustering taken as the one with the highest variance of information compared to the other clusterings, i.e. the most representative.

Each cluster produced by GPClust is associated with a defined mean and covariance function, which can be discretized to a multivariate normal distribution. In order to generate superclusters, the mean function was hierarchically clustered using correlation distance, and stability was assessed using pvclust (Suzuki and Shimodaira, 2006) with a 10,000 iterations. Groups of clusters found to be stable (approximately unbiased p-value > 0.95) were identified and defined as superclusters. Similar results were obtained using symmetric Kullback Liebler-divergence (also called relative entropy) as a distance metric. TI_50_ (time to 50% inhibition) for each cluster was calculated using the estimated mean function, by calculating the time taken to reach 50% of the initial estimated starting value.

#### Functional enrichment analysis

Gene Ontology (GO) enrichments performed using GOStats (Falcon and Gentleman, 2007) using all genes differentially expressed (FDR < 10%) as background. Terms with an FDR < 5% were defined as significantly enriched. Results from GO enrichments were simplified for presentation purposes by filtering terms with a high semantic similarity (Yu et al., 2010).

#### Transcription factor binding sites (TFBS) analysis

TFBS motifs were obtained from the JASPAR2016 database (Mathelier et al., 2016). Due to the impact of background sequence selection on the downstream results of motif identification (Roider et al., 2009; Worsley-Hunt et al., 2014), promoters (defined as +/− 500bp from the Ensembl annotated transcription start site (TSS)) were classified into different sets based on their GC content. Each set of promoters were then scanned using FIMO (Bailey et al., 2009) with a p-value threshold of 5×10^−4^ against the appropriate background. Enrichment of motifs in the promoter of genes in a specific cluster was calculated using a hypergeometric test and p-values were corrected using FDR after removing those motifs that were not present in the cluster of interest.

#### Comparison with transcriptional responses upon TCF7L2 KD

Microarray data was obtained from GEO:GSE48367 (Forrest et al., 2013) and processed using LIMMA (Linear Models for Microarray and RNA-Seq Data) (Ritchie et al., 2015). Normalized expression values were log_2_-transformed. Microarray probes that did not map to an annotated gene in Ensembl version 79 were removed from the analysis, and in the case of multiple probes mapping to one gene, the probe with the largest interquartile range was retained. Genes were defined as significantly downregulated using an FDR threshold of 10% and a fold-change of 0.5. Significance of the overlap between genes downregulated in SH-SY5Y following TCFKD and *Wnt-activated* genes was calculated using a hypergeometric with the number of genes expressed in both the HPAF-II time course and in the microarray experiment as background. Bonferroni correction was used to correct for multiple testing and control false positives.

#### Analysis of existing TCF7L2 ChIP-seq datasets

TCF7L2 peaks for various cell lines were obtained from ReMap 2018 (Chèneby et al., 2018; Frietze et al., 2012; Hazelett et al., 2014; Ni et al., 2013; Trompouki et al., 2011). The distance from a TCF7L2 peaks were assigned to their nearest gene promoter, defined as the Ensembl defined TSS, were calculated.

#### Proteomics

Tumors were homogenized on dry ice and solubilized with 8M urea and 20 mM HEPES. After reduction and alkylation, proteins were digested with trypsin overnight. Peptides in 0.1% TFA were separated using an Ultimate 3000 RSLC nano liquid chromatography system coupled to a Q-Exactive mass spectrometer (both Thermo Scientific) via an EASY-Spray source. Technical duplicates of 500 ng were loaded onto a trap column (Acclaim PepMap 100 C18, 100 μm, 2 cm) at 8 μl/min in 2 % acetonitrile, 0.1 % TFA. Peptides were eluted on-line to an analytical column (PepMap C18, 75 μm × 25 cm) and separated using a ramped 120 minute gradient, 445 % of buffer B (80 % acetonitrile, 0.1 % formic acid). Following data-dependent acquisition, raw data files were loaded into Progenesis QI (PQI v3, Non-linear Dynamics), which performed peak detection, chromatographic run alignment, combined peak list creation (Mascot mgf file) and protein quantitation. Search results (filtered to approximately 1% FDR) were imported back into Progenesis QI and relative protein quantitation was performed using non-conflicting peptides only. Protein normalization was performed by Progenesis QI (Ritchie et al., 2015).

## Supplementary Figures

**Supplemental Figure 1:**
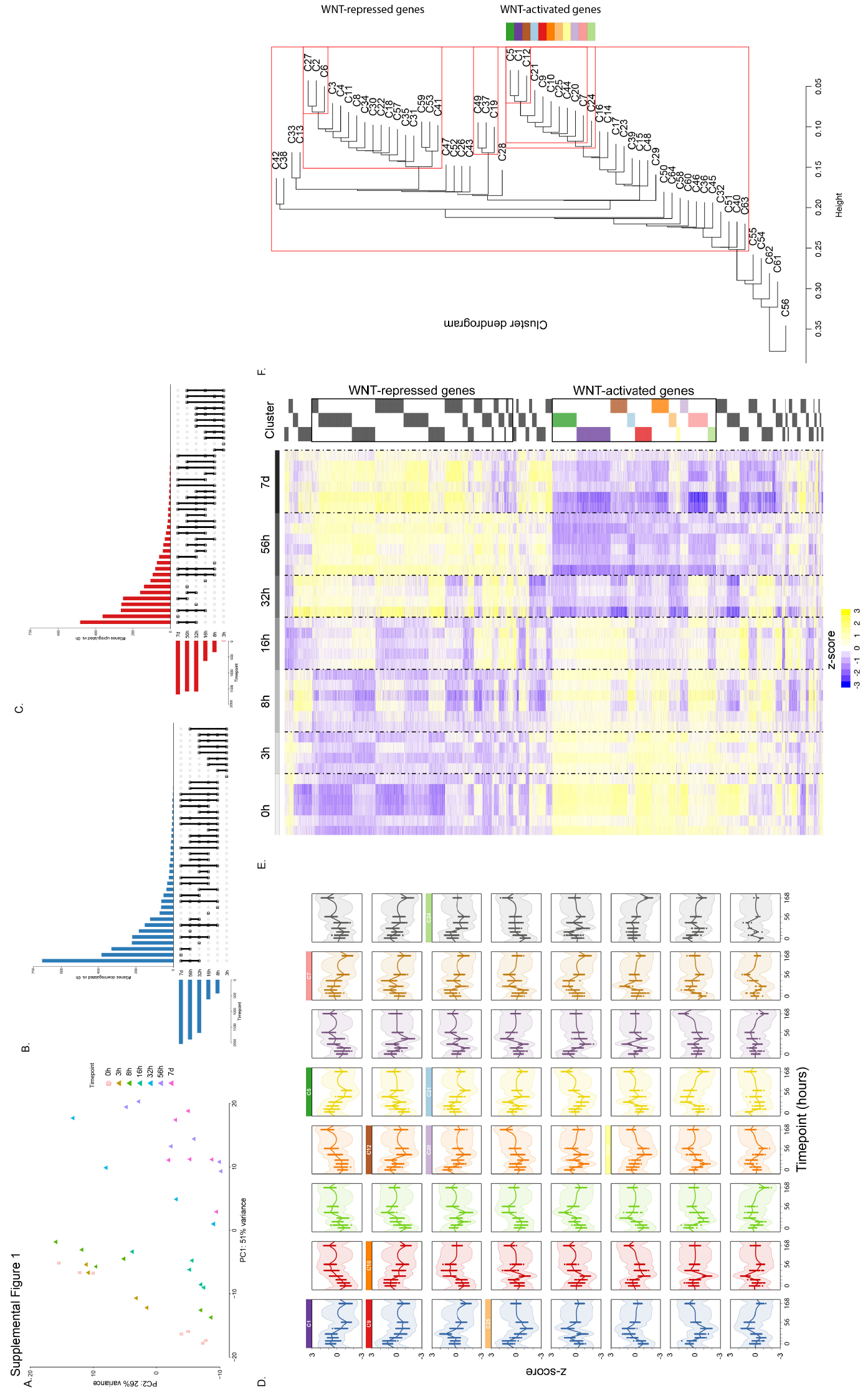
Time series transcriptomic analysis identifies clusters of genes with distinct dynamics in response to PORCN inhibition (accompanying Figure 1) A. Principal components analysis of HPAF-II orthotopic time course RNA-seq reveals coherent gene expression changes in response to ETC-159 over time.
B. Upset plot of downregulated genes reveals groups of genes that are downregulated across different sets of timepoints compared to 0 h (FDR < 10%, fold change < 0.5)
C. Upset plot of upregulated genes reveals groups of genes that are upregulated across different sets of timepoints compared to 0 h in response to PORCN inhibition (FDR < 10%, fold change > 1.5)
D. Mean and covariance of all clusters generated using GPclust. Clusters of genes defined as *Wnt-activated* are highlighted. Clusters are ordered by their size.
E. Heatmap displaying all genes that are differentially expressed over time (FDR < 10%), grouped into 64 distinct clusters based on their dynamics in response to PORCN inhibition.
F. Hierarchical clustering of the mean function of each cluster. Distinct stable groupings (i.e. super-clusters) were identified using pvclust with 1,000 iterations.

**Supplemental Figure 2:**
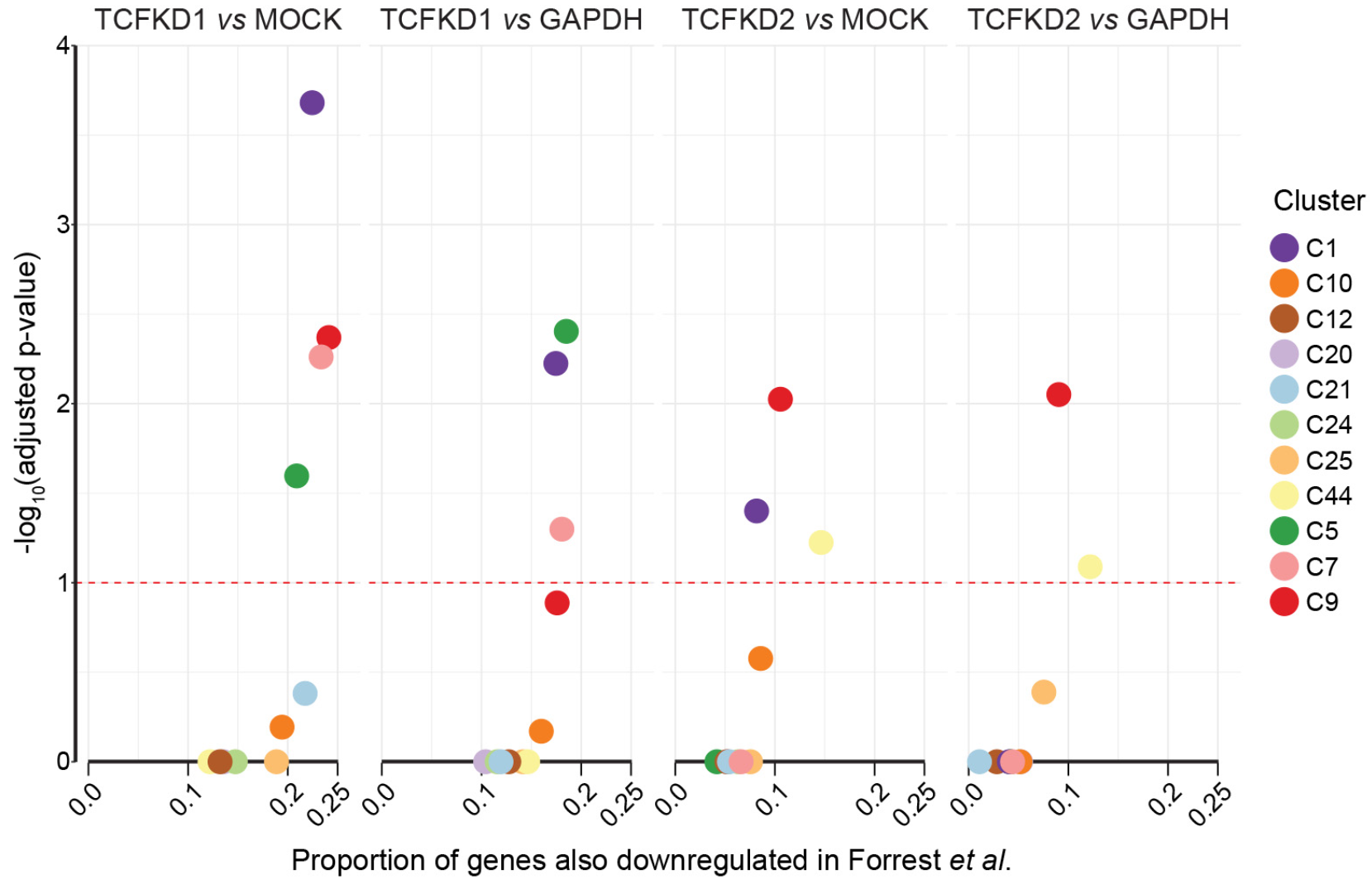
Clusters of WNT-activated genes are enriched for WNT targets identified following TCF KD in SH-SY5Y cells (accompanying Figure 1). The Forrest et al. dataset (Forrest et al., 2013) was reanalyzed (Methods) and the TCF-dependent genes identified. Examining the set of genes expressed in both datasets, we determined how many of our *Wnt-activated* genes were also downregulated in Forrest et al. (FDR < 10%, fold change < 0.5). The x-axis shows the proportion of genes in each Wnt-activated cluster that were also found to be downregulated following TCF knockdown. In no case did the cell line data contain more than 25% of the genes identified in our orthotopic model.

**Supplemental Figure 3:**
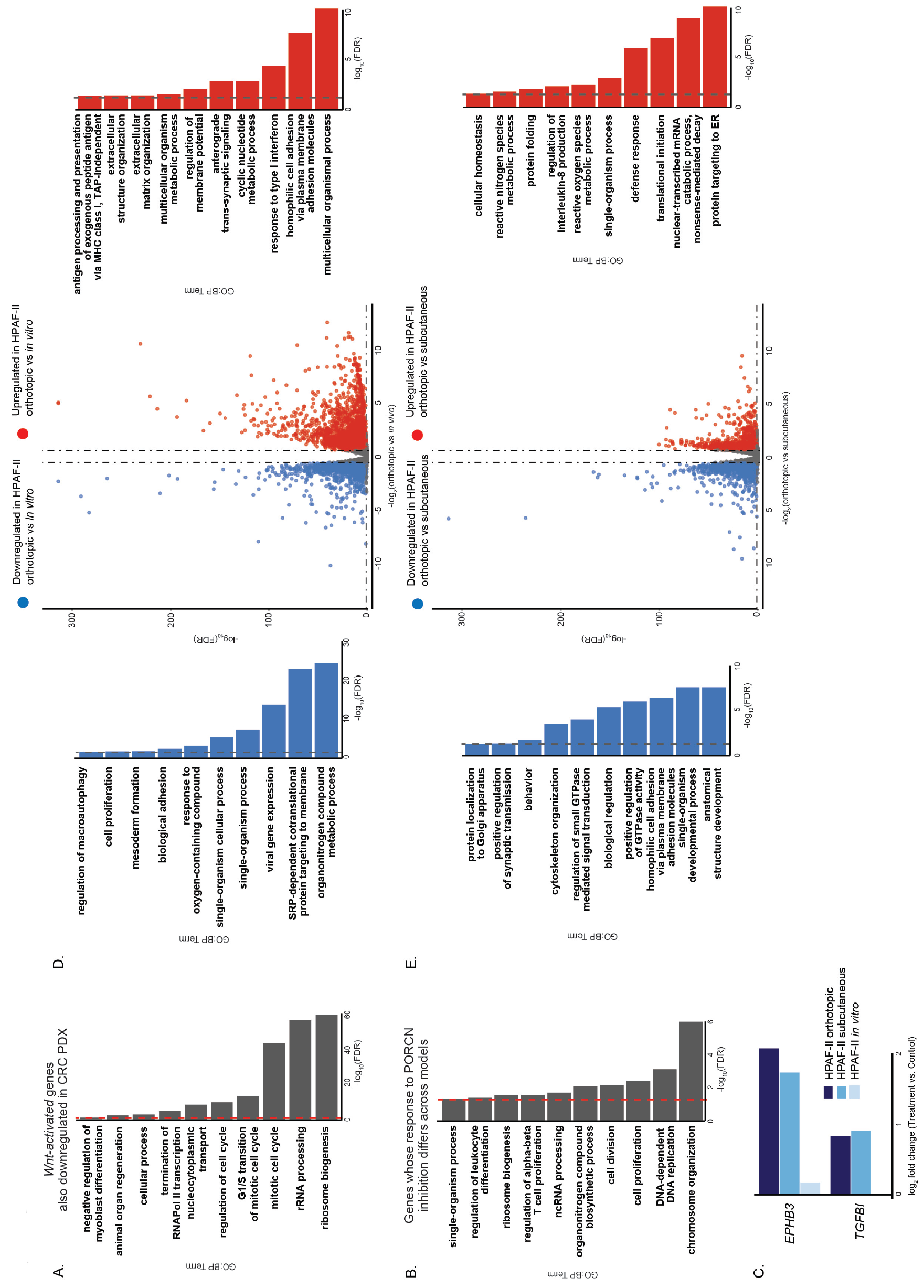
Comparison of models of Wnt-addicted cancer reveals genes and processes that differ in baseline expression and/or response to ETC-159 (accompanying Figure 2). A. *Wnt-activated* genes that are downregulated (FDR < 10%) in both orthotopic HPAF-II and CRC PDX models are enriched for cell cycle and ribosome biogenesis genes as assessed by GO:BP enrichment (FDR < 10%, dotted red line).
B. Genes whose response to PORCN inhibition differs across models (interaction test, FDR < 10%) are enriched for cell cycle processes as assessed by GO:BP enrichment (FDR < 5%, dotted red line).
C. *TGBI* and *EPHB3* do not respond *in vitro* but are WNT targets in both HPAF-II *in vivo* models.
D. Volcano plot and GO enrichments show that the genes expressed even before treatment differ widely between orthotopic and in vitro models (FDR < 10%). These genes are associated with distinct biological processes as assessed by GO:BP enrichment (FDR < 5%, dotted line).
E. Volcano plot and GO enrichments show that the genes expressed before treatment also differ between orthotopic and subcutaneous models (FDR < 10%). These genes are associated with distinct biological processes as assessed by GO:BP enrichment (FDR < 5%, dotted line).

**Supplemental Figure 4:**
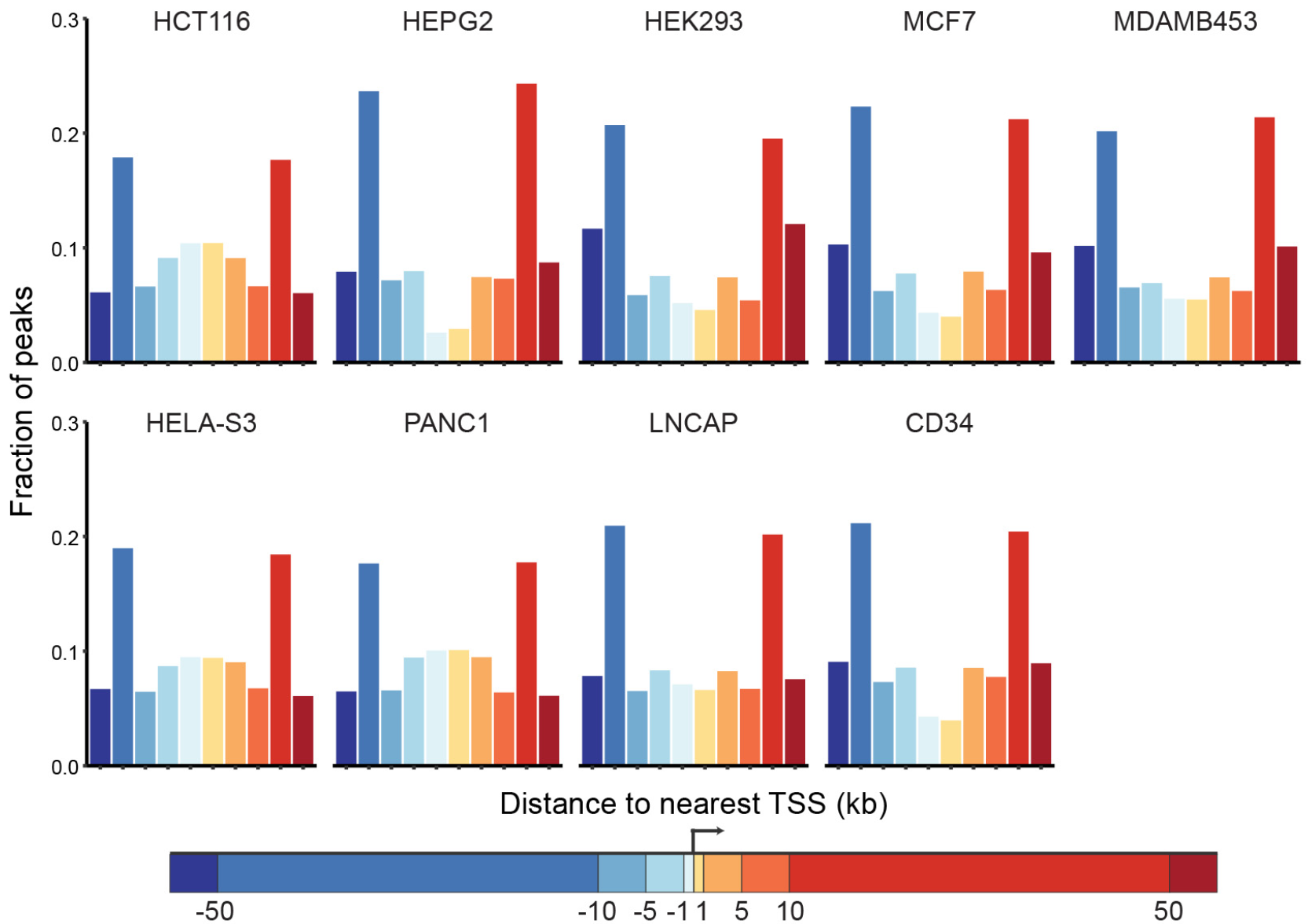
The majority of TCF7L2 peaks are not located near promoters (accompanying Figure 3). Analysis of existing TCF7L2 ChIP-seq data reveals that the majority of TCF7L2 binding peaks are located away (<5kb) from the promoters of genes.

**Supplemental Figure 5:**
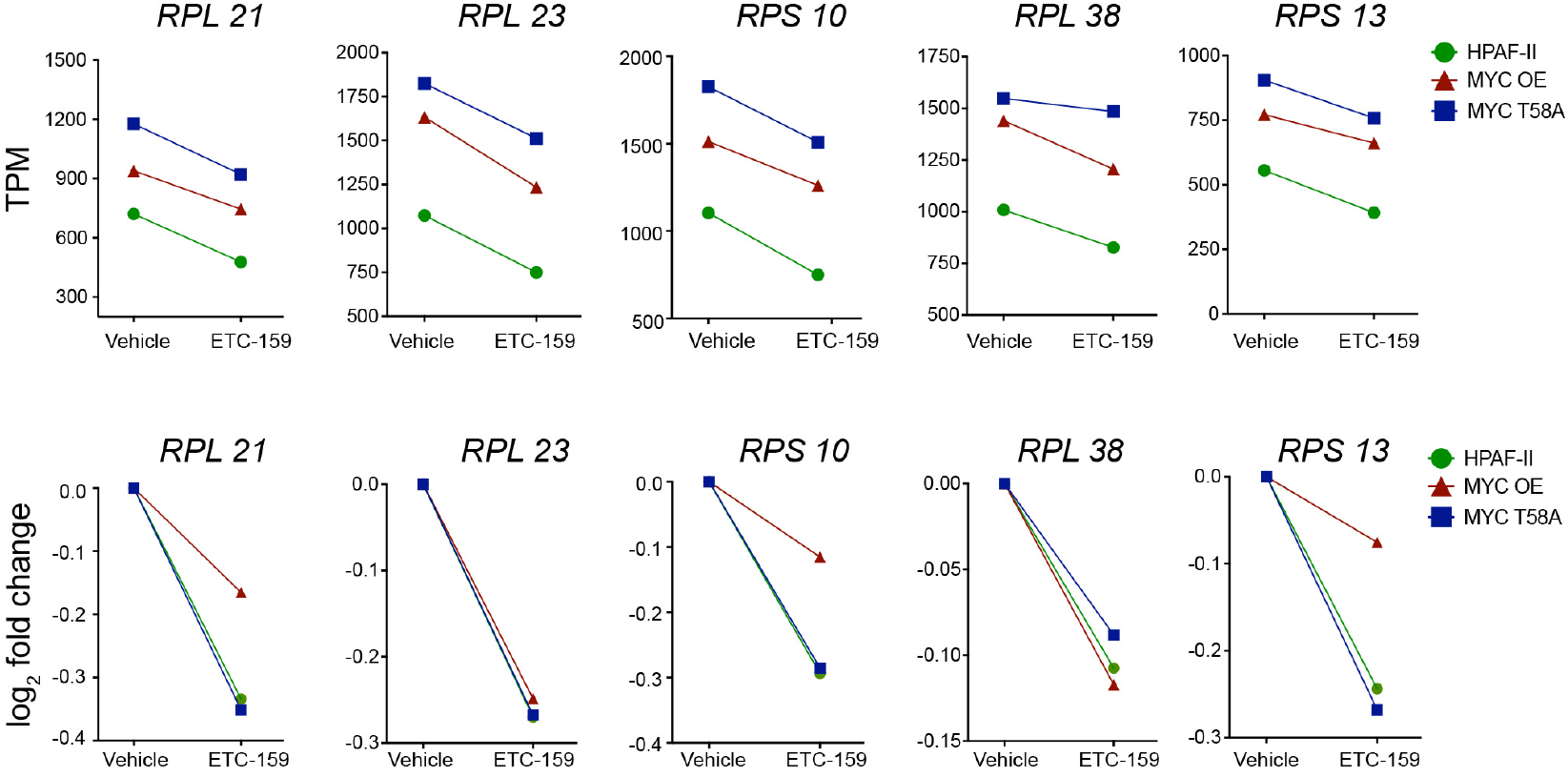
The response of RPLs and RPSs to PORCN inhibition is Myc-independent (accompanying Figure 7). Expression of representative RPL and RPS genes is enhanced by MYC and is dependent on Wnt signaling.

## References

Acebron, S.P., and Niehrs, C. (2016). β-catenin-Independent Roles of Wnt/LRP6 Signaling. Trends Cell Biol 26, 956–967.

Acebron, S.P., Karaulanov, E., Berger, B.S., Huang, Y.-L., and Niehrs, C. (2014). Mitotic Wnt signaling promotes protein stabilization and regulates cell size. Mol Cell 54, 663–674.

Annibali, D., Whitfield, J.R., Favuzzi, E., Jauset, T., Serrano, E., Cuartas, I., Redondo-Campos, S., Folch, G., Gonzàlez-Juncà, A., Sodir, N.M., et al. (2014). Myc inhibition is effective against glioma and reveals a role for Myc in proficient mitosis. Nat Commun 5, 4632.

Arnold, H.K., Zhang, X., Daniel, C.J., Tibbitts, D., Escamilla-Powers, J., Farrell, A., Tokarz, S., Morgan, C., and Sears, R.C. (2009). The Axin1 scaffold protein promotes formation of a degradation complex for c-Myc. Embo J 28, 500–512.

Byrne, A.T., Alférez, D.G., Amant, F., Annibali, D., Arribas, J., Biankin, A.V., Bruna, A., Budinská, E., Caldas, C., Chang, D.K., et al. (2017). Interrogating open issues in cancer precision medicine with patient-derived xenografts. Nat Rev Cancer 17, 254–268.

Cancer Genome Atlas Research Network (2017). Integrated Genomic Characterization of Pancreatic Ductal Adenocarcinoma. Cancer Cell 32, 185–203.e13.

Chen, B., Dodge, M.E., Tang, W., Ma, Z., Fan, C.-W., Wei, S., Williams, N.S., Roth, M.G., Amatruda, J.F., Chen, C., et al. (2009). Small molecule-mediated disruption of Wnt-dependent signaling in tissue regeneration and cancer. Nat Chem Biol 5, 100–107.

Chien, A.J., Conrad, W.H., and Moon, R.T. (2009). A Wnt survival guide: from flies to human disease. J Investig Dermatol 129, 1614–1627.

Coombs, G.S., Yu, J., Canning, C.A., Veltri, C.A., Covey, T.M., Cheong, J.K., Utomo, V., Banerjee, N., Zhang, Z.H., Jadulco, R.C., et al. (2010). WLS-dependent secretion of WNT3A requires Ser209 acylation and vacuolar acidification. J Cell Sci 123, 3357–3367.

Dang, C.V. (2010). Enigmatic MYC Conducts an Unfolding Systems Biology Symphony. Genes Cancer 1, 526–531.

Dang, C.V., Reddy, E.P., Shokat, K.M., and Soucek, L. (2017). Drugging the “undruggable” cancer targets. Nat Rev Cancer 17, 502–508.

Dolfini, D., and Mantovani, R. (2013). Targeting the Y/CCAAT box in cancer: YB-1 (YBX1) or NF-Y? Cell Death Differ 20, 676–685.

Finch, A.J., Soucek, L., Junttila, M.R., Swigart, L.B., and Evan, G.I. (2009). Acute overexpression of Myc in intestinal epithelium recapitulates some but not all the changes elicited by Wnt/beta-catenin pathway activation. Mol Cell Biol 29, 5306–5315.

Gabay, M., Li, Y., and Felsher, D.W. (2014). MYC activation is a hallmark of cancer initiation and maintenance. Cold Spring Harb Perspect Med 4, a014241–a014241.

Giraldez, A.J., Copley, R.R., and Cohen, S.M. (2002). HSPG modification by the secreted enzyme Notum shapes the Wingless morphogen gradient. Dev Cell 2, 667–676.

Hensman, J., Rattray, M., and Lawrence, N.D. (2015). Fast Nonparametric Clustering of Structured Time-Series. IEEE Trans Pattern Anal Mach Intell 37, 383–393.

Janda, C.Y., Waghray, D., Levin, A.M., Thomas, C., and Garcia, K.C. (2012). Structural Basis of Wnt Recognition by Frizzled. Science 337, 59–64.

Jiang, X., Hao, H.-X., Growney, J.D., Woolfenden, S., Bottiglio, C., Ng, N., Lu, B., Hsieh, M.H., Bagdasarian, L., Meyer, R., et al. (2013). Inactivating mutations of RNF43 confer Wnt dependency in pancreatic ductal adenocarcinoma. Proc Natl Acad Sci USA 110, 12649–12654.

Ju, X., Ishikawa, T.-O., Naka, K., Ito, K., Ito, Y., and Oshima, M. (2014). Context-dependent activation of Wnt signaling by tumor suppressor RUNX3 in gastric cancer cells. Cancer Sci 105, 418–424.

Killion, J.J., Radinsky, R., and Fidler, I.J. (1998). Orthotopic models are necessary to predict therapy of transplantable tumors in mice. Cancer Metastasis Rev 17, 279–284.

Koch, S., Acebron, S.P., Herbst, J., Hatiboglu, G., and Niehrs, C. (2015). Post-transcriptional Wnt Signaling Governs Epididymal Sperm Maturation. Cell 163, 1225–1236.

Kraushar, M.L., Viljetic, B., Wijeratne, H.R.S., Thompson, K., Jiao, X., Pike, J.W., Medvedeva, V., Groszer, M., Kiledjian, M., Hart, R.P., et al. (2015). Thalamic WNT3 Secretion Spatiotemporally Regulates the Neocortical Ribosome Signature and mRNA Translation to Specify Neocortical Cell Subtypes. J Neurosci 35, 10911–10926.

Liu, J., Pan, S., Hsieh, M.H., Ng, N., Sun, F., Wang, T., Kasibhatla, S., Schuller, A.G., Li, A.G., Cheng, D., et al. (2013). Targeting Wnt-driven cancer through the inhibition of Porcupine by LGK974. Proc Natl Acad Sci USA 110, 20224–20229.

Ly, L.L., Yoshida, H., and Yamaguchi, M. (2013). Nuclear transcription factor Y and its roles in cellular processes related to human disease. Am J Cancer Res 3, 339–346.

Madan, B., Ke, Z., Harmston, N., Ho, S.Y., Frois, A.O., Alam, J., Jeyaraj, D.A., Pendharkar, V., Ghosh, K., Virshup, I.H., et al. (2016). Wnt addiction of genetically defined cancers reversed by PORCN inhibition. Oncogene 35, 2197–2207.

Madan, B., and Virshup, D.M. (2015). Targeting Wnts at the source--new mechanisms, new biomarkers, new drugs. Mol. Cancer Ther. 14, 1087–1094.

Moon, R.T., and Gough, N.R. (2016). Beyond canonical: The Wnt and β-catenin story. Sci Signal 9, eg5–eg5.

Myant, K., and Sansom, O.J. (2011). Wnt/Myc interactions in intestinal cancer: partners in crime. Exp Cell Res 317, 2725–2731.

Nakamura, Y., de Paiva Alves, E., Veenstra, G.J.C., and Hoppler, S. (2016). Tissue- and stage-specific Wnt target gene expression is controlled subsequent to β-catenin recruitment to cis-regulatory modules. Development 143, 1914–1925.

Nusse, R., and Clevers, H. (2017). Wnt/β-catenin Signaling, Disease, and Emerging Therapeutic Modalities. Cell 169, 985–999.

Nusse, R., and Varmus, H. (2012). Three decades of Wnts: a personal perspective on how a scientific field developed. Embo J 31, 2670–2684.

Ong, C.K., Subimerb, C., Pairojkul, C., Myint, S.S., Rajasegaran, V., Wu, Y., Huang, D., Qian, C.-N., Ooi, A., Yongvanit, P., et al. (2012). Exome sequencing of liver fluke-associated cholangiocarcinoma. Nat Genet 44, 690–693.

Pelletier, J., Thomas, G., and Volarević, S. (2018). Ribosome biogenesis in cancer: new players and therapeutic avenues. Nat Rev Cancer 18, 51–63.

Pfister, A.S., and Kühl, M. (2018). Of Wnts and Ribosomes. Prog Mol Biol Transl Sci 153, 131–155.

Proffitt, K.D., Madan, B., Ke, Z., Pendharkar, V., Ding, L., Lee, M.A., Hannoush, R.N., and Virshup, D.M. (2013). Pharmacological inhibition of the Wnt acyltransferase PORCN prevents growth of WNT-driven mammary cancer. Cancer Res 73, 502–507.

Quin, J.E., Devlin, J.R., Cameron, D., Hannan, K.M., Pearson, R.B., and Hannan, R.D. (2014). Targeting the nucleolus for cancer intervention. Biochim Biophys Acta 1842, 802–816.

Ramakrishnan, A.-B., and Cadigan, K.M. (2017). Wnt target genes and where to find them. F1000Res 6, 746.

Reed, K.R., Athineos, D., Meniel, V.S., Wilkins, J.A., Ridgway, R.A., Burke, Z.D., Muncan, V., Clarke, A.R., and Sansom, O.J. (2008). B-catenin deficiency, but not Myc deletion, suppresses the immediate phenotypes of APC loss in the liver. Proc. Natl. Acad. Sci. U.S.a. 105, 18919–18923.

Rios-Esteves, J., and Resh, M.D. (2013). Stearoyl CoA Desaturase Is Required to Produce Active, Lipid-Modified Wnt Proteins. Cell Reports 4, 1072–1081.

Ruggero, D., and Pandolfi, P.P. (2003). Does the ribosome translate cancer? Nat Rev Cancer 3, 179–192.

Sansom, O.J., Meniel, V.S., Muncan, V., Muncan, V., Phesse, T.J., Wilkins, J.A., et al. (2007). Myc deletion rescues Apc deficiency in the small intestine. Nature 446, 676–679.

Sears, R., Nuckolls, F., Haura, E., Taya, Y., Tamai, K., and Nevins, J.R. (2000). Multiple Ras-dependent phosphorylation pathways regulate Myc protein stability. Genes Dev 14, 2501–2514.

Seshagiri, S., Stawiski, E.W., Durinck, S., Modrusan, Z., Storm, E.E., Conboy, C.B., Chaudhuri, S., Guan, Y., Janakiraman, V., Jaiswal, B.S., et al. (2012). Recurrent R-spondin fusions in colon cancer. Nature 488, 660–664.

Stevens, M.L., Chaturvedi, P., Rankin, S.A., Macdonald, M., Jagannathan, S., Yukawa, M., Barski, A., and Zorn, A.M. (2017). Genomic integration of Wnt/β-catenin and BMP/Smad1 signaling coordinates foregut and hindgut transcriptional programs. Development 144, 1283–1295.

Sulima, S.O., Hofman, I.J.F., De Keersmaecker, K., and Dinman, J.D. (2017). How Ribosomes Translate Cancer. Cancer Discov 7, 1069–1087.

Taelman, V.F., Dobrowolski, R., Plouhinec, J.-L., Fuentealba, L.C., Vorwald, P.P., Gumper, I., Sabatini, D.D., and De Robertis, E.M. (2010). Wnt Signaling Requires Sequestration of Glycogen Synthase Kinase 3 inside Multivesicular Endosomes. Cell 143, 1136–1148.

Takada, R., Satomi, Y., Kurata, T., Ueno, N., Norioka, S., Kondoh, H., Takao, T., and Takada, S. (2006). Monounsaturated fatty acid modification of Wnt protein: its role in Wnt secretion. Dev Cell 11, 791–801.

van den Heuvel, S., and Dyson, N.J. (2008). Conserved functions of the pRB and E2F families. Nat Rev Mol Cell Biol 9, 713–724.

van Riggelen, J., Yetil, A., and Felsher, D.W. (2010). MYC as a regulator of ribosome biogenesis and protein synthesis. Nat Rev Cancer 10, 301–309.

Wilkins, J.A., and Sansom, O.J. (2008). C-Myc is a critical mediator of the phenotypes of Apc loss in the intestine. Cancer Res 68, 4963–4966.

Yu, J. (2014). Updating the Wnt pathways. Biosci. Rep. 34, 593–607.

Zhang, X., Farrell, A.S., Daniel, C.J., Arnold, H., Scanlan, C., Laraway, B.J., Janghorban, M., Lum, L., Chen, D., Troxell, M., et al. (2012). Mechanistic insight into Myc stabilization in breast cancer involving aberrant Axin1 expression. Proc. Natl. Acad. Sci. U.S.a. 109, 2790–2795.

## Supplemental References

Bailey, T.L., Boden, M., Buske, F.A., Frith, M., Grant, C.E., Clementi, L., Ren, J., Li, W.W., and Noble, W.S. (2009). MEME SUITE: tools for motif discovery and searching. Nucleic Acids Res 37, W202–W208.

Chèneby, J., Gheorghe, M., Artufel, M., Mathelier, A., and Ballester, B. (2018). ReMap 2018: an updated atlas of regulatory regions from an integrative analysis of DNA-binding ChIP-seq experiments. Nucleic Acids Res 46, D267–D275.

Conway, T., Wazny, J., Bromage, A., Tymms, M., Sooraj, D., Williams, E.D., and Beresford-Smith, B. (2012). Xenome--a tool for classifying reads from xenograft samples. Bioinformatics 28, i172–i178.

Dobin, A., Davis, C.A., Schlesinger, F., Drenkow, J., Zaleski, C., Jha, S., Batut, P., Chaisson, M., and Gingeras, T.R. (2013). STAR: ultrafast universal RNA-seq aligner. Bioinformatics 29, 15–21.

Falcon, S., and Gentleman, R. (2007). Using GOstats to test gene lists for GO term association. Bioinformatics 23, 257–258.

Forrest, M.P., Waite, A.J., Martin-Rendon, E., and Blake, D.J. (2013). Knockdown of human TCF4 affects multiple signaling pathways involved in cell survival, epithelial to mesenchymal transition and neuronal differentiation. PLoS ONE 8, e73169.

Frietze, S., Wang, R., Yao, L., Tak, Y.G., Ye, Z., Gaddis, M., Witt, H., Farnham, P.J., and Jin, V.X. (2012). Cell type-specific binding patterns reveal that TCF7L2 can be tethered to the genome by association with GATA3. Genome Biol 13, R52.

Hazelett, D.J., Rhie, S.K., Gaddis, M., Yan, C., Lakeland, D.L., Coetzee, S.G., Ellipse/GAME-ON consortium, Practical consortium, Henderson, B.E., Noushmehr, H., et al. (2014). Comprehensive functional annotation of 77 prostate cancer risk loci. PLoS Genet 10, e1004102.

Li, B., and Dewey, C.N. (2011). RSEM: accurate transcript quantification from RNA-Seq data with or without a reference genome. BMC Bioinformatics 12, 323.

Love, M.I., Huber, W., and Anders, S. (2014). Moderated estimation of fold change and dispersion for RNA-seq data with DESeq2. Genome Biol 15, 550.

Mathelier, A., Fornes, O., Arenillas, D.J., Chen, C.-Y., Denay, G., Lee, J., Shi, W., Shyr, C., Tan, G., Worsley-Hunt, R., et al. (2016). JASPAR 2016: a major expansion and update of the open-access database of transcription factor binding profiles. Nucleic Acids Res 44, D110–D115.

Ni, M., Chen, Y., Fei, T., Li, D., Lim, E., Liu, X.S., and Brown, M. (2013). Amplitude modulation of androgen signaling by c-MYC. Genes Dev 27, 734–748.

Ritchie, M.E., Phipson, B., Wu, D., Hu, Y., Law, C.W., Shi, W., and Smyth, G.K. (2015). limma powers differential expression analyses for RNA-sequencing and microarray studies. Nucleic Acids Res 43, e47–e47.

Roider, H.G., Lenhard, B., Kanhere, A., Haas, S.A., and Vingron, M. (2009). CpG-depleted promoters harbor tissue-specific transcription factor binding signals--implications for motif overrepresentation analyses. Nucleic Acids Res 37, 6305–6315.

Storey, J.D., and Tibshirani, R. (2003). Statistical significance for genomewide studies. Proc Natl Acad Sci USA 100, 9440–9445.

Suzuki, R., and Shimodaira, H. (2006). Pvclust: an R package for assessing the uncertainty in hierarchical clustering. Bioinformatics 22, 1540–1542.

Trerè, D. (2000). AgNOR staining and quantification. Micron 31, 127–131.

Trompouki, E., Bowman, T.V., Lawton, L.N., Fan, Z.P., Wu, D.-C., DiBiase, A., Martin, C.S., Cech, J.N., Sessa, A.K., Leblanc, J.L., et al. (2011). Lineage regulators direct BMP and Wnt pathways to cell-specific programs during differentiation and regeneration. Cell 147, 577–589.

Worsley-Hunt, R., Mathelier, A., Del Peso, L., and Wasserman, W.W. (2014). Improving analysis of transcription factor binding sites within ChIP-Seq data based on topological motif enrichment. BMC Genomics 15, 472.

